# Direct binding of a fungal effector by the wheat RWT4 tandem kinase activates defense

**DOI:** 10.1101/2024.04.30.591956

**Authors:** Yi-Chang Sung, Yinghui Li, Zoe Bernasconi, Suji Baik, Soichiro Asuke, Beat Keller, Tzion Fahima, Gitta Coaker

## Abstract

Plants have intricate innate immune receptors that detect pathogens. Research has intensely focused on two receptor classes recognizing external and internal threats. Recent research has identified a class of disease resistance proteins called tandem kinase proteins (TKPs). We investigated RWT4, a wheat TKP that confers resistance to the devastating fungal pathogen *Magnaporthe oryzae*. We established a rice protoplast system, revealing RWT4 specifically recognizes the AvrPWT4 effector, leading to the transcription of defense genes and inducing cell death. RWT4 possesses both kinase and pseudokinase domains, with its kinase activity essential for defense. RWT4 directly interacts with and transphosphorylates AvrPWT4. Biolayer interferometry revealed both RWT4 kinase and pseudokinase regions bind the effector. Sequence similarity and structural modeling revealed an integrated partial kinase duplication in RWT4’s kinase region as critical for effector interaction and defense activation. Collectively, these findings demonstrate that TKPs can directly bind a recognized effector, leading to downstream defense activation.

## Introduction

Plant genomes encode multiple innate immune receptors capable of recognizing diverse pathogen classes ^1^. These immune receptors recognize pathogen features such as conserved molecular patterns or specialized secreted pathogen effector proteins ^1^. The main classes of plant immune receptors are broadly categorized into cell-surface localized receptors, including receptor-like kinases (RLKs) or receptor-like proteins (RLPs), as well as intracellular nucleotide-binding domain leucine-rich repeat (NLR) immune receptors ^2,3^. Most identified and well-studied receptors belong to RLKs/RLPs or NLRs, also known as canonical resistance genes ^4^. Recently, kinase fusion proteins (KFP) have emerged as new players in plant immunity. KFPs contain distinct protein architecture, including dual-kinase domains termed tandem kinase proteins (TKPs) or kinase fused with an integrated domain ^5^.

To date, more than 100 TKPs have been found across plant species, spanning both dicots and monocots ^6,7^. A more recent TKP atlas comprising 104 angiosperm genomes identified 2,682 TKPs^8^. TKPs represent a protein, not a gene, family since their kinase domains can be derived from different kinase (sub)families ^9^. However, TKPs conferring disease resistance have been exclusively identified from the *Triticeae* tribe. Currently, ten *Triticeae* TKPs have been identified that confer resistance to biotrophic or hemibiotrophic fungal pathogens ^5,9,10^. Barley RPG1 conferring resistance to barley stem rust was the first TKP identified ^11^. Subsequent cloning of wheat resistance proteins, WTK1 to WTK7-TM and RWT4, also revealed dual-kinase architecture ^6,10,12–15^. KFPs can also contain integrated non-kinase domains. For example, Sr43 (kinase-DUF347-DUF668) and WTK6-vWA (TKP-vWA), have been cloned with resistance to stem rust and leaf rust, respectively ^16,17^. NLR immune receptors can also carry integrated domains responsible for directly binding recognized pathogen effectors, leading to defense activation and cell death ^18^. However, it is unclear if TKPs can directly recognize pathogen effectors or other pathogen components.

Current data, primarily from mutant analyses, indicates that both domains are required for TKP-mediated resistance, which includes eliciting plant cell death. Wheat Tandem Kinase 1 (WTK1), conferring resistance to yellow rust, induces rapid and localized cell death post-haustoria formation, suggesting that defense activation requires the perception of an unknown effector ^9^. Mutagenesis assays have revealed the requirement of both kinase and pseudokinase domains for WTK1 function ^9^. Similarly, mutants in RPG1’s kinase or pseudokinase domain exhibited full susceptibility to *Puccinia graminis* f. sp. *tritici* ^19^. RPG1 exhibits *in vitro* autophosphorylation activity, is rapidly phosphorylated upon exposure to avirulent spores, and can also induce cell death ^19,20^. These data indicate that TKPs are rapidly activated upon pathogen perception and can elicit hallmarks of innate immune responses, including cell death ^6,10,14,21,22^.

Recently, the *Rwt4* TKP was cloned from the D-genome of hexaploid wheat ^15^. RWT4 recognizes the PWT4 effector from *Magnaporthe oryzae* (syn *Pyricularia oryzae*) pathotype *Avena*, leading to an incompatible interaction ^23^. *M. oryzae* is composed of host-specific subgroups that can cause disease in rice, finger millet, oat, perennial ryegrass, and wheat. *M. oryzae* poses a threat to global wheat production ^24^. The *M. oryzae* host jump to wheat required loss of pathogen recognition by the TKP *Rwt4* and the NLR *Rwt3* ^15^.

Here, we examined RWT4 as a model for TKP activation. We demonstrate that RWT4 directly binds and phosphorylates the recognized effector AvrPWT4, but not an unrecognized VirPWT4 allele. Furthermore, RWT4 can be transferred to a rice protoplast system and confer defense activation in the presence of AvrPWT4. Structure-function analyses identified key RWT4 residues required for effector binding and defense activation. These findings provide detailed mechanistic insights into TKP effector binding and activation.

## Results

### RWT4 recognizes AvrPWT4 in wheat and rice

*Rwt4* is located in the D genome of hexaploid wheat ^15^. First, we phenotyped the hexaploid wheat cultivars Norin 4 (*Rwt4*), Cadenza (*Rwt4*), Hope (*rwt4*), and Chinese Spring (*rwt4*) for their responses to inoculation with spores of *M. oryzae* pathotype *Triticum* (MoT) isolate Br48 and its *AvrPwt4* transformants. Disease symptoms were recorded four days post-inoculation (dpi). Leaves infected with Br48 showed moderate to severe symptoms across all tested samples. As expected, the Br48 transformant carrying *AvrPwt4* failed to cause comparable symptoms in Norin 4 and Cadenza, demonstrating that lines harboring *Rwt4* are resistant to *AvrPwt4*-carrying strains (Fig. 1a, Table S1). All genotypes we tested lack *Rmg8*, an MCTP-kinase that recognizes another *M. oryzae* effector ^25,26^.

**Figure 1.**
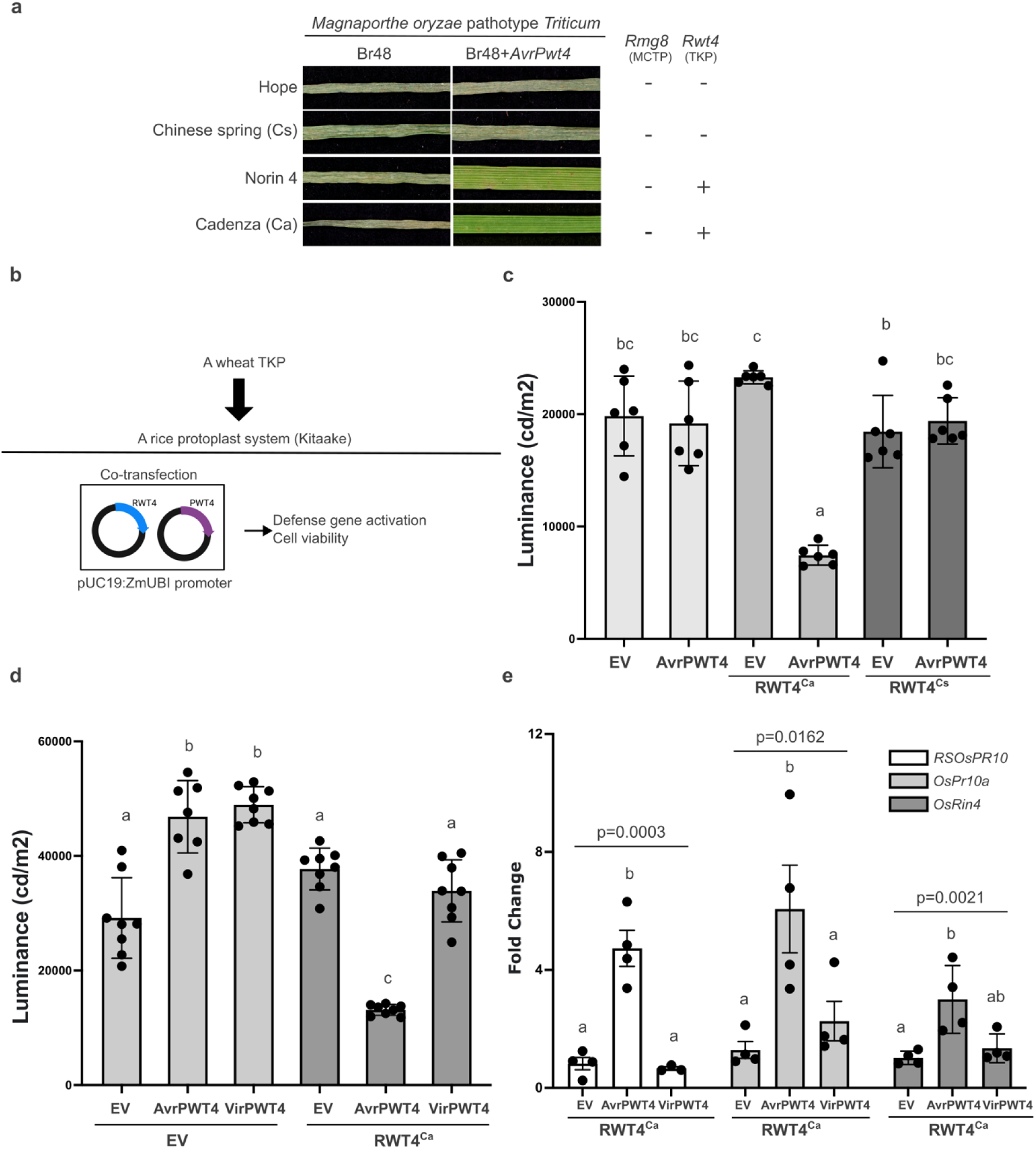
RWT4-mediated resistance requires AvrPWT4 perception. **(a)** Disease phenotyping of wheat cultivars Hope, Chinese Spring (Cs), Norin 4, and Cadenza (Ca) with *Magnaporthe oryzae* pathotype *Triticum* Br48 and a Br48 transformant carrying *AvrPwt4* (Br48+ *AvrPwt4*). Wheat genotypes with (+) or without (-) functional *Rmg8* or *Rwt4* alleles are indicated. **(b)** Experimental procedure of a rice protoplast transfection system for cell viability assays and defense gene expression tests. Genes of interest (*Rwt4* or *Pwt4*) were cloned to a pUC19 vector containing a maize polyubiquitin-1 (*Zm*Ubi-1) promoter. **(c)** RWT4 perceives AvrPWT4 triggering cell death responses in rice protoplasts. RWT4 from Cadenza (Ca) and Chinese Spring (Cs) were assayed for their perception of AvrPWT4 in rice protoplasts. Cell viability was measured as ATP activity presented in luminance (cd/m^2^). Luminescence (cd/m^2^) was measured 16 h after protoplast transfection. Six independent protoplast transfection events were used as biological replicates. Mean±SEM are presented. Differences between treatments were analyzed using one-way ANOVA with post-hoc Tukey HSD, p<0.05 **(d)** Only the recognized AvrPWT4 effector can induce RWT4^Ca^-triggered cell death. *Rwt4^Ca^* was co-transfected with either the avirulent (*AvrPwt4*) or the virulent (*VirPwt4*) allele of *Pwt4* in rice protoplasts. Protoplasts transfected with empty vector (EV) or co-transfected with empty vector and *Pwt4* were treated as experimental controls. Experiments were performed and analyzed as described in (c). **(e)** RWT4^Ca^ recognition of AvrPWT4 activates defense gene expression. Expression fold change of *RSOsPR10* (Os12g36830), *OsPr10a* (Os12g36850) and *OsRin4* (Os04g0379600) defense genes were measured 7 h after protoplast transfection. Gene expression was normalized to the internal control *OsUbq5* (Os01g22490), and data were normalized to the samples transfected with *Rwt4^Ca^*+EV as expression fold changes. Mean±SEM from four transfection events is presented. Differences between treatments were analyzed using one-way ANOVA with post-hoc Tukey HSD, p<0.05.

Next, we sought to identify a rapid plant system to investigate TKP activation. We employed a rice protoplast system to transiently co-express *Rwt4* and *Pwt4* and assayed defense responses, including cell death and defense gene expression (Fig. 1b). Luminance-based cell viability assays were conducted to evaluate the ability of *Rwt4* to specifically recognize *AvrPwt4* when expressed in rice. Protoplasts co-expressing *AvrPwt4* and the resistant Cadenza allele of *Rwt4* (*Rwt4^Ca^*) exhibited a significant reduction in viability compared to cells transfected with *Rwt4^Ca^* plus an empty vector (Fig. 1c, p<0.0001). In contrast, co-transfecting the susceptible Chinese Spring allele of *Rwt4* (*Rwt4^Cs^*) and *AvrPwt4* failed to cause cell death (Fig. 1c). Furthermore, we examined RWT4’s specificity in recognizing PWT4 for defense activation by co-transfecting *Rwt4^Ca^* with either *AvrPwt4* or *VirPwt4* in rice protoplasts. A significant strong reduction of cell viability was specifically observed in cells co-transfected with *Rwt4^Ca^* and *AvrPwt4* but not in cells co-transfected with either *VirPwt4* or an empty vector (Fig. 1d, p<0.0001). Similarly, we measured defense gene activation and observed that the defense marker genes *RSOsPR10*, *OsPr10a* and *OsRin4* ^27–29^ were specifically induced in cells co-transfected with *Rwt4^Ca^* and *AvrPwt4* (Fig. 1e). RWT4 and PWT4 effectors were detectable in rice protoplasts by western blot (Fig. S1). These results demonstrate that RWT4 is functional when transferred to rice and can specifically recognize the AvrPWT4 effector.

### The kinase activity of RWT4 is required for defense activation

*Rwt4* was identified as a tandem kinase encoding a kinase-pseudokinase domain organization. To explore the kinase domain family/subfamily of RWT4^Ca^, we conducted a sequence homology search for the RWT4^Ca^ kinase domain (amino acids 189-470) and pseudokinase domain (amino acids 554-840) in The Arabidopsis Information Resource (TAIR, www.arabidopsis.org), identifying AT4G00960 and AT4G05200 as kinase and pseudokinase orthologs, respectively. Analysis of their kinase family/subfamily using the annotated Arabidopsis kinome dataset ^30^ revealed that both of the domains belong to the LRR_8B kinase subfamily, predominantly associated with cysteine-rich receptor-like kinases for defense regulation. The two regions are interconnected by a linker region, which is flexible and intrinsically disordered (Fig. 2a). Except for WTK1, all other identified TKPs controlling disease resistance contain LRR_8B kinase domains ^9,10,16^.

**Figure 2.**
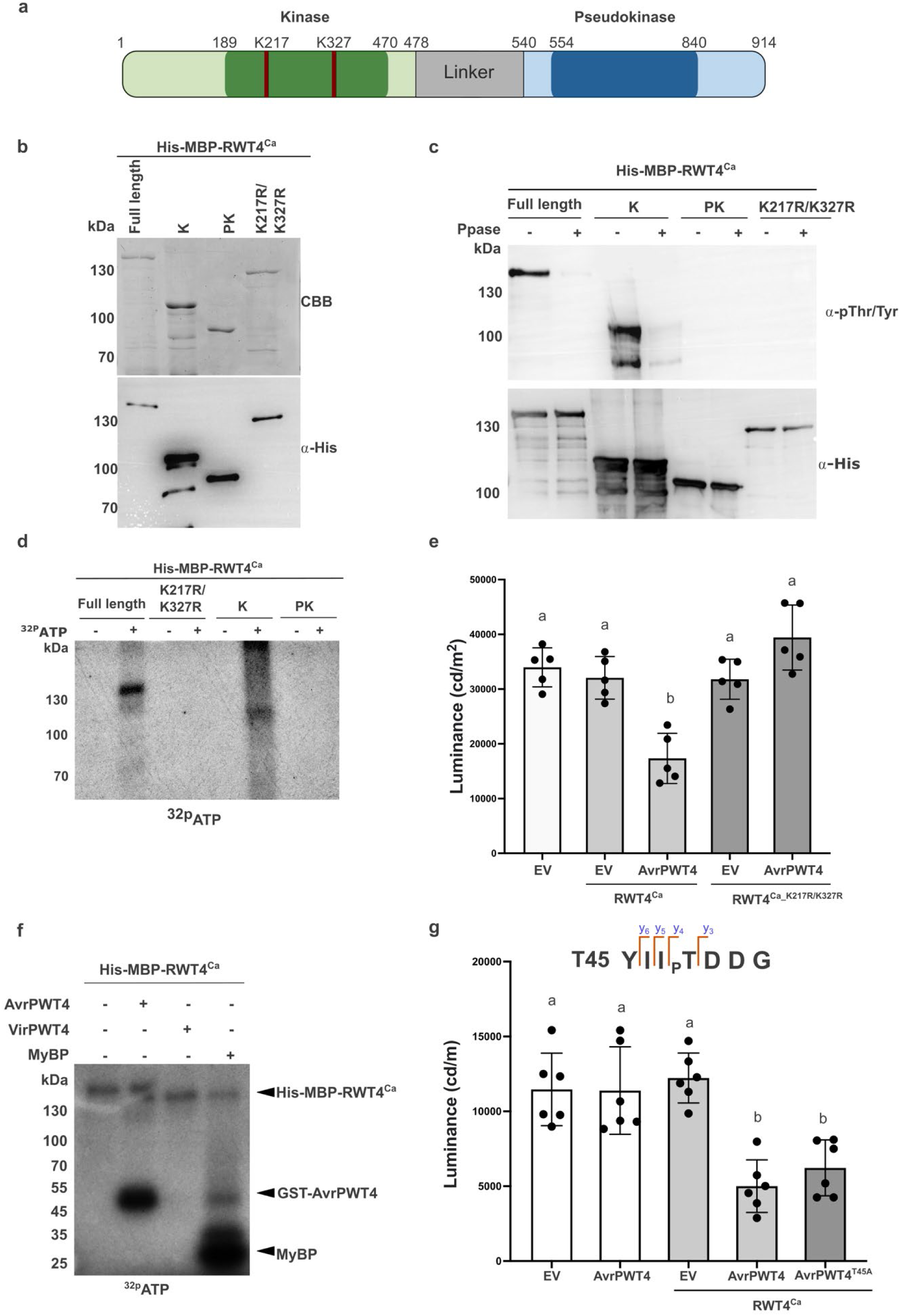
RWT4 is an active kinase and transphosphorylates AvrPWT4. **(a)** RWT4^Ca^ encodes kinase and pseudokinase architecture with a flexible linker between the two. The kinase and pseudokinase domain borders are indicated in dark green and dark blue, respectively. Predicted kinase active sites: K217 and K327. **(b)** Purity of recombinant proteins isolated from *E. coli.* His-MBP-RWT4^Ca^, kinase (His-MBP-K, amino acids 1-478), pseudokinase (His-MBP-PK, amino acids 540-914), and its kinase dead variant (RWT4^Ca_K217R/K327R^) are shown by Coomassie brilliant blue (CBB) stained SDS-PAGE and corresponding western blot using anti-His. **(c)** Autophosphorylation activity of His-MBP-RWT4^Ca^. Autophosphorylation status of the His-MBP-RWT4^Ca^ proteins were assayed by western blot using anti-phospho-Threonine/Tyrosine antibody (α-pThr/Tyr) with or without lambda protein phosphatase incubation prior to separating by SDS-PAGE. Anti-His western blotting (α-His) demonstrates equal loading of the His-MBP-RWT4^Ca^ proteins. **(d)** Active protein phosphorylation was measured by incubating the His-MBP-CaRWT4 proteins with [γ^-^ ^32P^]ATP before separating by SDS-PAGE and visualizing by autoradiography. **(e)** RWT4^Ca^-AvrPWT4 triggered cell death requires RWT4^Ca^ kinase activity. *Rwt4^Ca^* and the kinase dead variant (*Rwt4^Ca^*^_K217R/K327R^) were co-transfected with *AvrPwt4* into rice protoplasts for cell viability assays. Five independent protoplast transfection events were used as biological replicates and Mean±SEM are presented. Treatment differences were analyzed using one-way ANOVA with post-hoc Tukey HSD, p<0.05. **(f)** RWT4^Ca^ specifically trans-phosphorylates AvrPWT4. Transphosphorylation activity was assayed by mixing His-MBP-RWT4^Ca^ and AvrPWT4 at room temperature for 20 min before incubated with [γ-^32^P]ATP. Phosphorylation of the substrate proteins (AvrPWT4 or MyBP) were visualized via autoradiography. **(g)** Phosphorylated peptides were mapped to the sequence of AvrPWT4 for functional analysis. The phosphorylated peptide YIITDDG was identified from LC-MS/MS with the phosphorylation site T45. *Rwt4^Ca^* was co-transfected with either wild-type *AvrPwt4* or the phosphorylation null mutant *AvrPwt4^T45A^* to rice protoplasts. Luminescence (cd/m^2^) was measured 16 h after protoplast transfection as an indicator of cell viability. Six independent protoplast transfection events were used as biological replicates. Mean±SEM are presented. Differences between treatments were analyzed using one-way ANOVA with post-hoc Tukey HSD, p<0.05.

We explored the importance of kinase activity in RWT4-mediated defense. We expressed and purified recombinant RWT4 proteins including full length wild type RWT4^Ca^ (His-MBP-RWT4^Ca^), its kinase region (His-MBP-K), pseudokinase region (His-MBP-PK), and a kinase-dead variant (His-MBP-RWT4^Ca_K217R/K327R^), with mutations in the two conserved lysines located within the ATP binding motif (K217R) and the catalytic residue (K327R) predicted via InterPro ^31,32^ (Fig. 2b). Immunoblot analysis employing a phosphor-Thr/Tyr antibody detected phosphorylated His-MBP-RWT4^Ca^ and His-MBP-K while the signal was removed by treatment with lambda protein phosphatase (Fig. 2c), suggesting RWT4^Ca^ and His-MBP-K are capable of autophosphorylation. In contrast, RWT4’s pseudokinase region and MBP-RWT4^Ca_K217R/K327R^ exhibited no detectable phosphorylation by western blot (Fig. 2c). Subsequent kinase assays using [^32^P]γATP detected *de novo* phosphorylation of His-MBP-RWT4^Ca^ and His-MBP-K, whereas His-MBP-PK and His-MBP-RWT4^Ca_K217R/K327R^ failed to exhibit kinase activity (Fig. 2d). Western blot with anti-HIS and Coomassie-stained SDS-PAGE analysis further revealed a size shift between His-MBP-RWT4^Ca^ and His-MBP-RWT4^Ca_K217R/K327R^ (Fig. 2b and 2c), further confirming that RWT4^Ca_K217R/K327R^ serves as a kinase dead variant of RWT4^Ca^. These data demonstrate that RWT4 is an active kinase and contains kinase-pseudokinase architecture.

Next, we employed mass spectrometry to identify unique RWT4 phosphorylation sites upon incubation with AvrPWT4 *in vitro*. RWT4 was phosphorylated at multiple sites in the absence of AvrPWT4. Four unique phosphorylated residues were identified only after incubation with recombinant AvrPWT4, two located within the kinase and two within the pseudokinase (Fig. S2, S3). One of these phosphorylated residues, Y204, resides within the ATP binding motif (195-217 amino acids), just proximal to the glycine-rich loop of RWT4’s kinase domain (KD) and is a conserved residue in the kinase of other TKP LRR_8B members (Fig. S3). The remaining three residues, T164, S609, S803, are not conserved across the kinase/pseudokinase of other TKP LRR_8B members.

To determine if RWT4’s kinase activity is required for defense induction, we co-transfected *AvrPwt4* with either *Rwt4^Ca^* or *Rwt4^Ca_K217R/K327R^* and measured cell viability as an indicator of defense activation. Unlike wild type *Rwt4^Ca^*, which strongly reduced cell viability when recognizing *AvrPwt4*, cells co-transfected with *AvrPwt4* and *Rwt4^Ca_K217R/K327R^*exhibited no significant decrease in cell viability (Fig. 2e, p<0.0001). Western blot detected protein equal expression of RWT4^Ca^ and RWT4^Ca_K217R/K327R^ in rice protoplasts (Fig. S1a). These data collectively demonstrate that RWT4 kinase activity is necessary for defense responses mediated by perception of AvrPWT4.

### RWT4 phosphorylates AvrPWT4, but not the unrecognized effector VirPWT4

Next, we sought to investigate whether RWT4 phosphorylates AvrPWT4. Purified recombinant His-MBP-RWT4^Ca^ was mixed with GST-AvrPWT4 or GST-VirPWT4 at 1:1 molar ratio 30 min prior to the kinase assay using [^32^P]γATP. The common kinase substrate myelin basic protein (MyBP) was included as a positive control. Phosphorylated GST-AvrPWT4 was detected from samples incubated with His-MBP-RWT4^Ca^ (Fig. 2f) or His-MBP-K (Fig. S4a). However, no phosphorylated GST-AvrPWT4 was observed from samples incubated with His-MBP-PK (Fig. S4a) or the kinase dead variant His-MBP-RWT4^Ca_K217R/K327R^ (Fig. S4b). Furthermore, RWT4^Ca^ did not phosphorylate GST-VirPWT4 when using recombinant proteins (Fig. 2f). These findings demonstrate that RWT4^Ca^specifically transphosphorylates the recognized effector AvrPWT4 *in vitro*.

To further investigate the biological significance of AvrPWT4 phosphorylation, we mapped AvrPWT4 phosphorylation sites by mass-spectrometry and identified five phosphorylated residues (Fig. S5a). Comparison of recognized and unrecognized alleles of PWT4 revealed that the T45 residue is unique to AvrPWT4 but absent in VirPWT4 (Fig. S5b, Fig. 2g). We generated the phosphorylation null mutant *AvrPwt4^T45A^* and tested its ability to be recognized by RWT4^Ca^ in rice protoplasts. However, *AvrPwt4^T45A^* had a similar ability to induce cell death in protoplasts as wild-type *AvrPwt4* when co-transfected with *Rwt4^Ca^*, suggesting the effector phosphorylation at this residue is not required for RWT4-mediated defense (Fig. 2g, p<0.0001). Western blot detected AvrPWT4 and AvrPWT4^T45A^ expression from protoplasts (Fig. S5c).

### RWT4^Ca^ directly and specifically interacts with AvrPWT4

It has been an unsolved question if TKPs can act as a receptor for a corresponding effector. Yeast two-hybrid demonstrated an interaction between RWT4^Ca^ and AvrPWT4 but no interaction with VirPWT4 (Fig. 3a). Western blot analysis confirmed the protein expression of RWT4^Ca^, AvrPWT4, and VirPWT4 in yeast (Fig. S6). To further validate that the RWT4 TKP can directly interact with AvrPWT4, we utilized Bio-layer interferometry (BLI) to characterize binding kinetics. RWT4^Ca^ was purified with a His-tag for immobilizing to the biosensors, while AvrPWT4 and VirPWT4 were expressed with a GST tag which was subsequently removed by TEV protease treatment during purification (Fig. 3b). Based on the molecular weight and yields of purified proteins, a 1.5x dilution of molar concentration gradient of effectors were prepared from 10 µM to 51 nM as ligands, while purified His-RWT4^Ca^ was immobilized to the Octet NTA biosensors as analytes. Strong binding kinetics between RWT4^Ca^ were observed for all tested concentrations of AvrPWT4 showing wavelength shifts from 0.6844 nm to 0.0264 nm, with binding constants (*K_D_*) less than 1x10^-12^ (Fig. 3c and Table S2). In contrast, the binding affinity of RWT4^Ca^ to VirPWT4 is weak and unstable, showing wavelength shifts from 0.1754 nm to 0.0023 nm with a *K_D_* value greater than 10^-7^ (Fig. 3c and Table S2). Background binding to GST exhibited similar weak kinetics comparable to the binding observed between His-RWT4^Ca^ and VirPWT4 (Fig. 3c and Table S2). These results demonstrate that RWT4^Ca^ has direct, stable, and specific binding to AvrPWT4.

**Figure 3.**
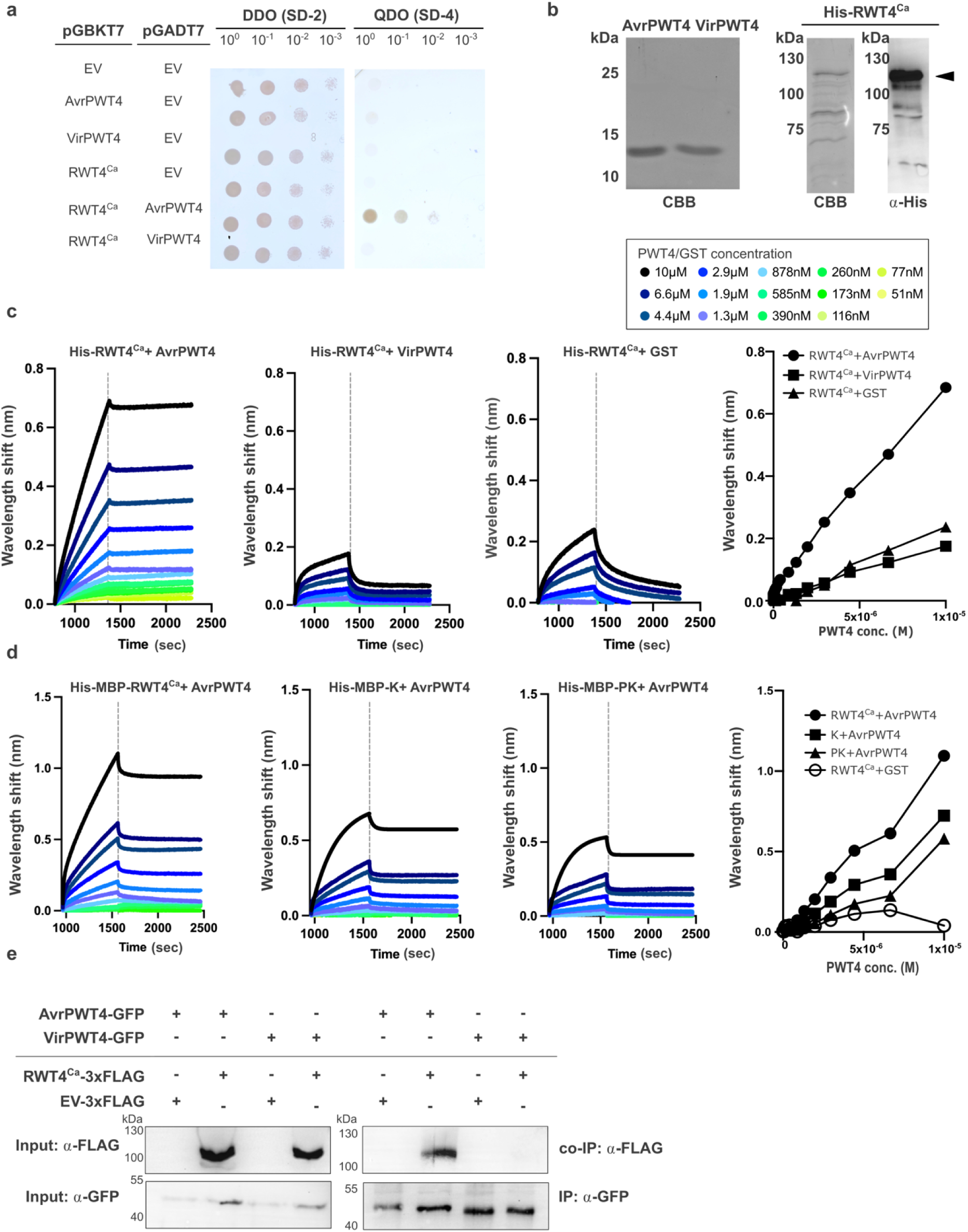
RWT4^Ca^ directly and specifically binds AvrPWT4. **(a)** Yeast-two hybrid (Y2H) assays using *Rwt4^Ca^*, *AvrPwt4* and *VirPwt4*. *Rwt4^Ca^* was cloned to the bait vector pGBKT7, while *AvrPwt4* and *VirPwt4* were cloned into the prey vector pGADT7. Yeast growth on DDO (SD-Leu/-Trp) media indicates the presence of the two plasmids, and the growth on QDO/X/A (SD-Ade/-His/-Leu/-Trp/3-amino-1,2,4-triazole) media indicates interaction. Serial dilutions from cell suspensions of a single yeast colony expressing the bait and prey plasmids are shown and represent the strength of interaction. Empty vectors (EV) of pGBKT7 and pGADT7 were used as negative controls. **(b)** Purity of AvrPWT4, VirPWT4, and His-RWT4^Ca^. AvrPWT4 and VirPWT4 were expressed and purified with a GST tag that was removed and separated by TEV protease and gel filtration. Purity of His-CaRWT4 is shown by Coomassie brilliant blue (CBB) stained SDS-PAGE and Anti-His western blotting (α-His). **(c)** and **(d)** Bio-layer interferometry (BLI) characterizes binding affinity between His-RWT4^Ca^ and PWT4 proteins. **(c)** His-RWT4^Ca^ was mobilized to Ni-NTA biosensors as analytes and incubated with a range of concentrations (51 nM to 10 µM) of AvrPWT4, VirPWT4, or GST ligand proteins. Sensorgrams show BLI traces (wavelength shifts) during association and dissociation steps (shown as time at the x-axis) normalized to no-ligand controls. The steady state analysis (right) shows wavelength shifts against different ligand concentrations (AvrPWT4, VirPWT4, or GST, from 10 µM to 116 nM). **(d)** His-MBP-RWT4^Ca^, His-MBP-Kinase (K, amino acids 1-478) and His-MBP-Pseudokinase (PK, amino acids 540-914) proteins were immobilized to Ni-NTA biosensors and incubated with AvrPWT4. Binding affinity between His-MBP-RWT4^Ca^, His-MBP-K and His-MBP-PK to AvrPWT4 were detected by BLI during association and dissociation steps (shown as time at the x-axis) and presented in the steady state analysis against different concentrations of AvrPWT4 (right). **(e)** *In-planta* interaction of RWT4^Ca^ and AvrPWT4. AvrPWT4-GFP or VirPWT4-GFP were co-expressed with RWT4^Ca^-3xFLAG in *N. benthamiana* using *Agrobacterium*-mediated transient expression and subjected to anti-GFP immunoprecipitation (IP). Input and IP samples were immunoblotted with anti-FLAG and anti-GFP antibodies.

Further BLI assays were conducted to determine which RWT4 ^Ca^ domain is required for AvrPWT4 interaction. Due to challenges in obtaining high-quality proteins of kinase and pseudokinase with a His-tag, we expressed proteins with a His-MBP tag to promote solubility. Interestingly, both His-MBP-K and His-MBP-PK demonstrated measurable binding to AvrPWT4 within concentrations ranging from 10 µM to 173 nM, exhibiting similar binding constants although His-MBP-PK displayed weaker wavelength shifts (Fig. 3d and Table S3). Notably, the full-length His-MBP-RWT4^Ca^ exhibited stronger binding affinity than its individual domains, and all three tested proteins showed no binding to GST (Fig. 3d and Table S3).

To validate the association between RWT4^Ca^ and AvrPWT4, we performed co-immunoprecipitation assays in *Nicotiana benthamiana*. AvrPWT4-GFP or VirPWT4-GFP were co-expressed with RWT4^Ca^-3xFLAG or an empty vector (EV-3xFLAG). Immunoblotting results demonstrated that AvrPWT4-GFP strongly associated with RWT4^Ca^-3xFLAG, while no detectable interaction between VirPWT4-GFP and RWT4^Ca^-3xFLAG was observed (Fig. 3e). RWT4^Ca^-3xFLAG, AvrPWT4-GFP, and VirPWT4-GFP were expressed and detected by immunoblotting in *N. benthamiana* (Fig. 3e). Collectively, these *in vitro* and *in vivo* results indicate that RWT4 functions as a receptor for AvrPWT4, with both kinase and pseudokinase involved in effector binding.

### Structure-guided identification of RWT4 specificity

We further investigated the sequence or structural features determining the binding of RWT4 to AvrPWT4. Through pairwise sequence alignment, the N-terminus of RWT4 (amino acids 1-142) was found to exhibit similarity to another section (amino acids 150-290) comprising part of the kinase domain. InterPro annotations identified a partial kinase (amino acids 25-145) within the segment. Subsequent sequence alignment between this partial kinase and RWT4’ kinase domain identified amino acids 39-142 that exhibited 63.83% sequence similarity to a fragment of the kinase domain (amino acids 150-290) (Fig. 4a). This conserved region exhibits both DNA and protein sequence similarity to the KD and thus termed an integrated kinase duplication (KDup) (Fig. 4a and 4b). KDup contains a portion of a kinase ATP-binding motif but the predicted active site K217 is not included. By employing AlphaFold2, the structure model of RWT4^Ca^ revealed that the KDup (yellow) comprises a short α-helix and a β-sheet, resembling the N-terminus of the kinase domain (green) (Fig. 4c and Fig. S7a). These observations demonstrate RWT4^Ca^ contains an integrated partial kinase duplication, KDup.

**Figure 4.**
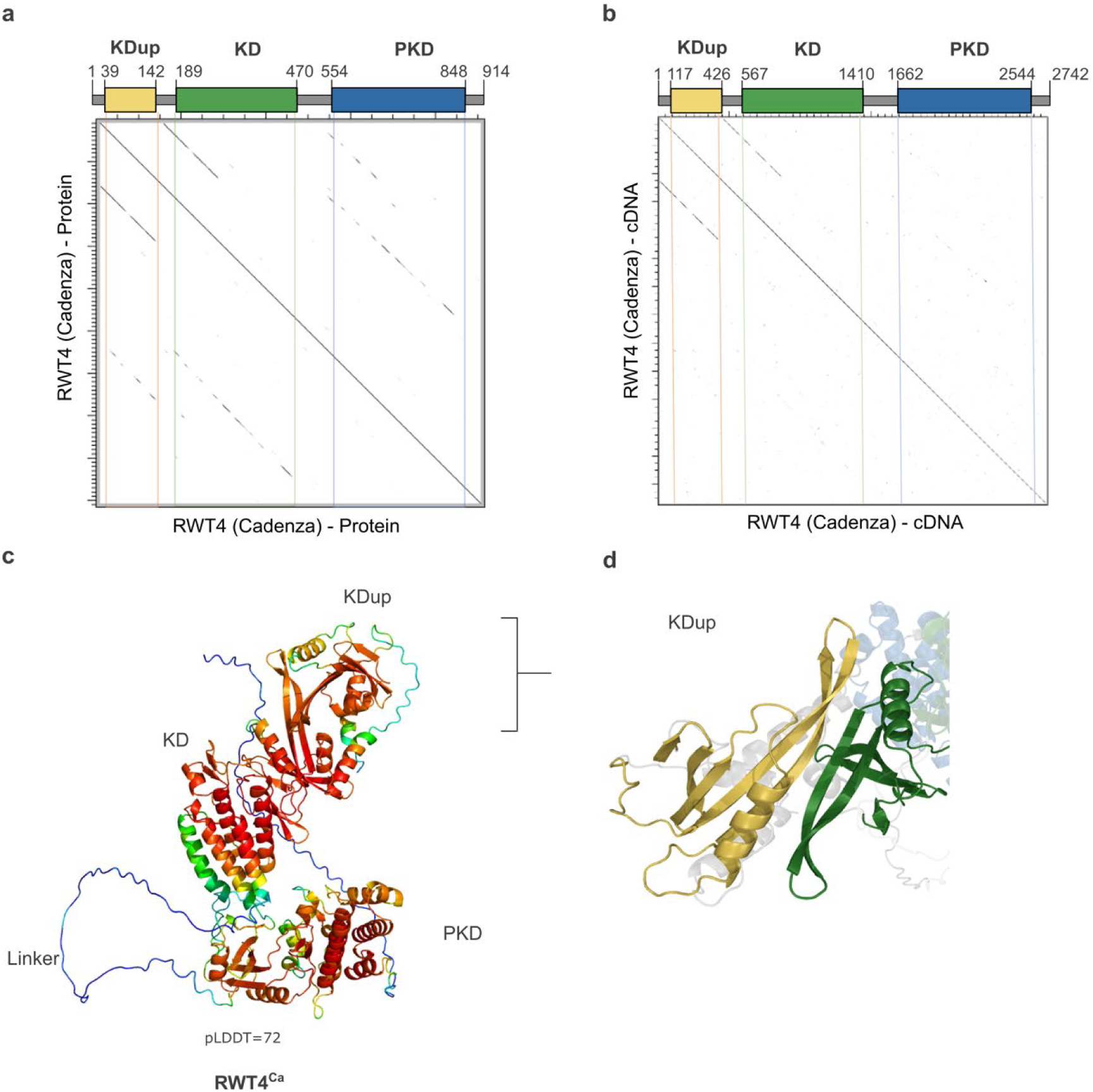
The kinase region of RWT4^Ca^ contains an integrated N-terminal truncated kinase duplication (KDup). **(a)** Dot plot of the RWT4^Ca^ protein demonstrating collinearity between KDup (amino acids 39-142) and a segment of the kinase domain in amino acids 150-290. KD = kinase domain, PKD = pseudokinase domain. Vertical lines denote domain borders, diagonal lines indicate collinearity. **(b)** Dot plot of the *RWT4^Ca^* cDNA demonstrating collinearity between KDup and a segment of the kinase domain (88.73% identity). **(c)** AlphaFold model of RWT4^Ca^, colored by pLDDT score. **(d)** Structural similarity between the KDup (yellow) and KD (green).

Next, we predicted the RWT4^Ca^-AvrPWT4 protein complex through AlphaFold-Multimer ^33^ (Fig. 5a, left and Fig. S7). Structural modeling suggests that AvrPWT4 (purple) mediates interactions between the KDup (yellow) and pseudokinase (blue). We identified potential binding regions in the KDup (117-127), the KD (261-270) and two regions in the PKD (551-560, 627-645). Next, we compared *Rwt4* alleles from known *MoT*-resistant wheat cultivars (Norin 4, Cadenza, Jagger, Paragon and Claire) with susceptible cultivars (Chinese Spring and CDC Stanley) ^15,23^ (Fig. S8 and Table S4). We identified sequence polymorphisms within the predicted KDup binding region (117-127) and PKD region 2 (627-645) (Fig. 5b). Additionally, a six-amino-acid extension at the C-terminus was specifically conserved in susceptible *Rwt4* alleles (Fig. 5b).

**Figure 5.**
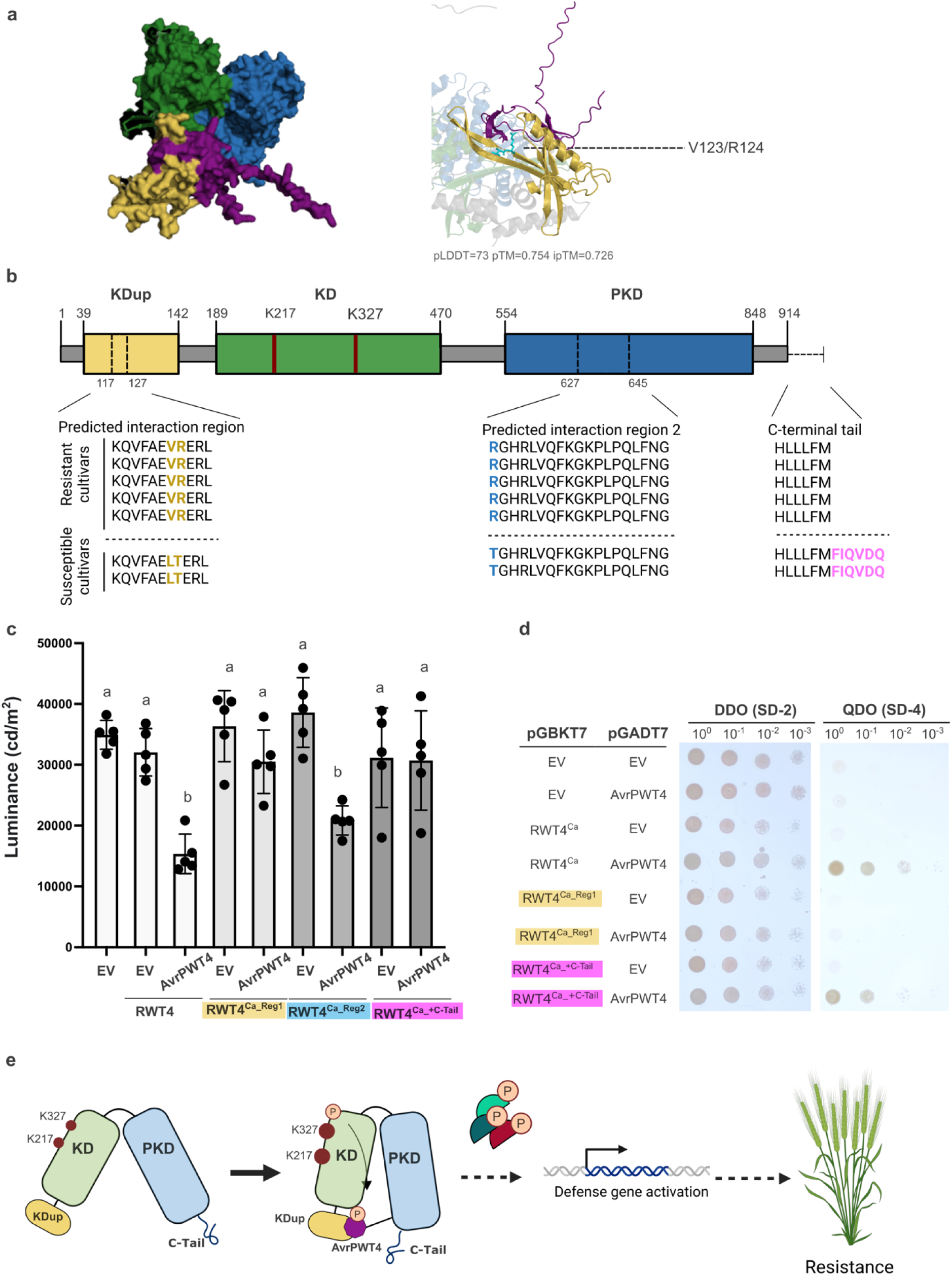
Structure-guided identification of RWT4 specificity. **(a)** AlphaFold Multimer modeling predicts that AvrPWT4 (purple) forms an interaction complex between the integrated kinase duplication (KDup, yellow) and pseudokinase (blue). The predicted binding region V123/R124 in RWT4’s KDup is highlighted (cyan, right panel). pLDDT indicates confidence in structure modeling and pTM and ipTM indicate the confidence for interaction. The kinase (green, 1-470) and pseudokinase (554-914) regions were separately inputted for modeling, with the linker region removed. **(b)** Illustration of sequence and structure features of RWT4^Ca^. *Rwt4* sequences were obtained from resistant wheat cultivars (Norin 4, Cadenza, Jagger, Paragon, and Claire) and susceptible wheat cultivars (Chinese Spring and CDC Stanley). Kinase active sites are indicated in red. The interaction regions were identified based on the modeling from (**a**). **(c)** RWT4^Ca^ variants validate the requirement of Region 1 and C-terminal sequences for defense activation. *Rwt4^Ca^* and its variants (*Rwt4*^Ca_Reg1^: V123L/R124T, *Rwt4*^Ca_Reg2^: R627T, *Rwt4*^Ca_+C-Tail^: RWT4^Ca^ with the extended C-terminal residues FIQVDQ) were co-transfected with *AvrPwt4* into rice protoplasts. Cell viability was measured as ATP activity presented in luminance (cd/m^2^). Data are represented as the mean ± SEM (n= 5). Different letters indicate significant differences using one-way ANOVA with post-hoc Tukey HSD (*P* ≤ 0.05). **(d)** Yeast-two hybrid confirms the interaction between RWT4^Ca^ variants to AvrPWT4. *Rwt4^Ca^* and its variants *Rwt4*^Ca_Reg1^, and *Rwt4*^Ca_+C-Tail^ were cloned to the bait vector pGBKT7 while *AvrPwt4* was cloned to the prey vector pGADT7. The transformation of bait and prey vectors were confirmed by yeast growth on DDO (SD-Leu/-Trp) media. The interaction was assayed by growth on QDO (SD-Ade/-His/-Leu/-Trp/3-amino-1,2,4-triazole) media. Serial dilutions from cell suspensions of a single yeast colony expressing the bait and prey plasmids are shown and represent the strength of interaction. Empty vectors (EV) of pGBKT7 and pGADT7 were used as negative controls. **(e)** RWT4 working model. KD = kinase domain, PKD = pseudokinase domain, KDup = integrated kinase duplication, P = phosphorylation, red = RWT4^Ca^ kinase active sites, C-Tail = C-terminal tail of RWT4.

To test the importance of the identified binding regions from AlphaFold-Multimer and sequence analyses, we generated RWT4 variants by replacing sequences between resistant and susceptible alleles in the context of RWT4^Ca^ (RWT4^Ca_Reg^^1^: V123L/R124T, RWT4^Ca_Reg^^2^: R627T) (Fig. 5b). A RWT4 variant incorporating the six-amino-acid extension at its C-terminus was also generated (RWT4^Ca_+C-tail^) (Fig. 5b). Subsequently, we evaluated the capacity of these RWT4 variants to perceive AvrPWT4 for defense activation in rice protoplasts. Protoplasts transfected with wild-type *Rwt4^Ca^* and *AvrPwt4* displayed a significant reduction in cell viability (Fig. 5c). However, cells transfected with the *Rwt4^Ca_Reg1^* variant and *AvrPwt4* failed to show a decrease in viability (Fig. 5c). Similarly, no cell death was observed upon co-transfection of *Rwt4^Ca_+C-tail^* with *AvrPwt4* (Fig. 5c). Cells expressing *Rwt4^Ca_Reg2^* and *AvrPwt4* exhibited a comparable reduction in cell viability to cells expressing *Rwt4^Ca^* and *AvrPwt4* (Fig. 4C, p<0.0001). Western blotting confirmed the protein expression of RWT4^Ca^, RWT4^Ca_Reg1^, RWT4^Ca_Reg2^, and RWT4^Ca_C-tail^ in rice protoplasts (Fig. S9a). These findings highlight the importance of RWT4^Ca^’s C-terminus and residues V124/R125 in the KDup in defense activation (Fig. 5a, right).

To assess the impact of these residues on AvrPWT4 binding, yeast-two hybrid analyses was conducted. As expected, RWT4^Ca^ can interact with AvrPWT4 (Fig. 5d and S8b). However, a weak interaction was observed between the susceptible variant from Chinese Spring, RWT4^Cs^, and AvrPWT4 (Fig. S9b). Next, we analyzed the interaction between RWT4 variant proteins and AvrPWT4. No interaction was observed between RWT4^Ca_Reg1^ and AvrPWT4, indicating that V124/R125 in the integrated KDup is required for AvrPWT4 binding (Fig. 5d). The C-terminal extension did not influence binding to AvrPWT4 (Fig. 5d). Western blot assays confirmed expression of RWT4 and its variants in yeast (Fig. S9c). These data demonstrate that the binding capacity of RWT4 to AvrPWT4 is required, but not sufficient for eliciting defense.

## Discussion

TKPs have emerged as a novel class of non-canonical resistance proteins, representing ∼10% of the cloned *Triticeae* R genes ^4,34^. Currently, TKPs conferring disease-resistance are exclusively found in monocots ^6^. In this study, we demonstrate the RWT4 TKP can be transferred and elicit defense between wheat and rice. Furthermore, both RWT4 kinase activity and direct effector binding are required for defense activation. It was hypothesized that TKP pseudokinase domains acts as a decoy for effector binding, activating the kinase domain for downstream phosphorylation or signaling ^9^. Our results demonstrate that AvrPWT4 binds to both kinase and pseudokinase regions. AvrPWT4 binding to both regions *in planta* may facilitate RWT4 intramolecular kinase interactions for activation (Fig. 5e). For plant receptor-like kinases, hetero-dimerization is a common activation mechanism for signaling initiation ^35^.

TKP-mediated resistance is associated with the induction of cell death and robust defense responses ^6,14,22^. Like PRR and NLR immune receptors that can directly detect pathogen components, RWT4 directly binds to AvrPWT4 and this binding is crucial for defense activation. *Rwt4* is allelic to *Wtk3*, which confers resistance to powdery mildew disease caused by *Blumeria graminis* f.sp. *tritici* ^14,15^. Though highly similar in amino acid sequence, the effector targets of WTK3 are currently unknown, and no ortholog of *Pwt4* is present in *B. graminis*. The majority of functional TKPs conferring disease resistance belong to the LRR_8B kinase subfamily ^9,10,16^. Prototypic members of this subfamily are cysteine-rich receptor-like kinases involved in plant defense and can confer resistance to fungal pathogens ^36,37^.

Whether an NLR is involved in TKP signaling downstream of TKP-effector binding remains an open question. Arabidopsis ZED1, a pseudokinase belonging to receptor-like cytoplasmic kinase (RLCK), targets the *P. syringae* effector HopZ1^38^. This interaction is crucial for HopZ1 recognition by the NLR ZAR1, leading to the formation of a resistance complex that activates immunity^38^. PBS1, another Arabidopsis RLCK, acts as a decoy that activates the NLR PBS5 upon cleavage by the *P. syringae* effector AvrPphB^39^. Although our results demonstrate that the direct binding of RWT4 to AvrPWT4 and RWT4 kinase activity are essential for eliciting immunity, the involvement of an NLR in TKP-mediated defense cannot be ruled out. RWT4 retains effector recognition when transferred between wheat and rice, indicating that an NLR may be ancient. The elucidation of TKP immune complexes and direct downstream targets will reveal how these novel proteins confer resistance.

*In vitro* kinase activity has been demonstrated for RPG1, WTK7-TM, and our work on the RWT4 TKP ^10,19^. *In vitro*, the pseudokinase region does not inhibit the kinase activity of RPG1 and RWT4 ^19^. We hypothesize that kinase activity is inhibited at a resting state *in planta* in the absence of pathogen perception. RPG1 activation *in planta* is only observed in the presence of avirulent spores of *P. graminis* ^20^, consistent with the hypothesis that TKPs rely on pathogen perception for activation. Additional regions are also required for RWT4 full functionality such as the V123R124 sites for effector binding and the C-terminus (Fig. 4c and 4d).

The identified effector binding site V123R124 is within the integrated kinase duplication, consistent with the hypothesis that KDup is an integrated domain (ID) for effector binding (Fig. 4 and 5b). IDs commonly found in NLRs include kinase, WRKY, Zinc-finger BED, and heavy metal-associated (HMA) domains^18^. IDs facilitate NLRs to recognize sequence/structure-diverse effectors that target to similar host proteins ^40^. The HMA domain of rice Pik-1 directly binds to the effector Avr-Pik1 for immune activation and the capability of effector binding contributes to positive selection for Pik-1 alleles ^41^. Similarly, in a companion paper Reveguk et al., (2024) recently identified IDs are prevalent in TKPs found across the plant kingdom ^8^. They also identified an integrated HMA domain in the first TKP to be cloned, barley RPG1^8^. No other TKPs conferring resistance aside from RWT4/WKT3 contain IDs that are partial kinase duplications. Thus, TKPs conferring disease resistance may frequently employ diverse IDs to trap pathogen effectors.

Effector targeting of plant kinases is an important strategy to enhance pathogen virulence. The *Pseudomonas syringae* AvrPtoB effector targets the plant kinase SnRK2.8, promoting pathogen virulence upon SnRK2.8 phosphorylation ^42^. The fungal NIS1 effector family from *Colletotrichum* spp. and *Magnaporthe oryzae* act as kinase inhibitors, targeting the BAK1 and BIK1 kinases required for effective defense mediated by multiple surface-localized receptors ^43^. PWT4 has canonical fungal effector features, including an N-terminal secretion signal and small size (75 amino acids) ^23^. AvrPWT4 suppresses resistance conferred by RMG8, an MCTP kinase fused with multiple C2 domains ^25,44^. Thus, AvrPWT4 likely targets plant kinases to promote virulence in the absence of RWT4. Accordingly, we hypothesize that KFPs serve both as receptors and decoys for perception and deception of biotrophic fungal effectors that are aiming to suppress host kinases involved in the activation of plant immunity ^5^.

TKPs have emerged as a novel class of resistance proteins that have been widely deployed to control devastating biotrophic fungal pathogens ^6,10,11,14,15^. Future investigations identifying effector-TKP pairs as well as their phosphorylation targets will shed light into TKP activation and signaling. This foundational information can facilitate receptor engineering and the deployment of TKPs across diverse plant species.

## Online Methods

### Wheat infection assays

Wheat seeds were pre-germinated on moistened filter papers. After 24 hours, sprouted seeds were sown in the soil in seedling cases (5.5 × 15 × 10 cm, Sakata Prime Mix, Sakata Seed Corporation, Yokohama, Japan), and grown at 22°C in a controlled-environment room with a 12-h photoperiod of fluorescent lighting for eight days. Primary leaves of the nine-day-old seedlings were fixed onto a hard plastic board with rubber bands just before inoculation. Conidial suspensions (1 × 10^5^ conidia/mL) were prepared as described previously ^45^ and sprayed onto fixed primary leaves using an air compressor. The inoculated seedlings were incubated in a tray sealed with plastic wrap under dark and humid conditions at 22°C for 24 h, then transferred to dry conditions with a 12-h photoperiod of fluorescent lighting and incubated at 22°C for an additional 3-5 days. Four to six days after inoculation, symptoms were evaluated based on the size and color of lesions ^46^. The size of lesions was rated on six progressive grades from 0-5: 0 = no visible infection; 1 = pinhead spots; 2 = small lesions (<1.5 mm); 3 = scattered lesions of intermediate size (<3 mm); 4 = large typical lesion; and 5 = complete blighting of leaf blades. A disease score comprised a number denoting the lesion size and a letter indicating the lesion color: ‘B’ for brown lesions and ‘G’ for green lesions. The infection types 0, 1B, and 2B were regarded as resistant, while the infection types 3G, 4G, and 5G were considered susceptible (Table S1).

### Cloning and site-directed mutagenesis

The *Rwt4* Cadenza allele (*Rwt4^Ca^*, TraesCAD_scaffold_040753_01G000200.1) and Chinese Spring allele (*Rwt4^Cs^*, TraesCS1D02G058900) were codon-optimized for expression in rice and synthesized (Twist Bioscience gene fragment). *Rwt4^Ca^*^_K217R/K327R^ was generated by PCR-based site-directed mutagenesis. Genes of interest were cloned to Gateway™ pENTR™ 4 Dual Selection Vector (A10465) by NEBuilder HiFi DNA Assembly Master Mix (NEB, E2621L) and cloned to a pUC19 expression vector powered by the maize ubiquitin promoter (*Zm*UBI) via Gateway LR reaction. Full length of *Rwt4^Ca^*, its kinase region, and pseudokinase region were cloned into *E. coli* expression plasmids pET28a-His-MBP ^47^ or pET28a-His (EMD Biosciences) using ligation-independent cloning (LIC) for recombinant protein production ^48^.

We refer to the *Pwt4* effector in two terms, the avirulent allele that can be recognized by *Rwt4* (*AvrPwt4*) and a virulent form that cannot be recognized (*VirPwt4*) ^23^. *AvrPwt4* (*Pwt4-Br58* type allele, GeneBank:LC202655.1) and *VirPwt4* (*Pwt4-Br48* type allele, GenBank: LC202656.1), lacking their N-terminal signal peptide as predicted by SignalP 5.0 (https://services.healthtech.dtu.dk/services/SignalP-5.0/), were codon optimized, synthesized (Twist Bioscience gene fragment) and cloned into the pUC19:*Zm*UBI expression vector via Gateway LR reaction. *AvrPwt4* and *VirPwt4* were cloned into an *E. coli* expression plasmid pET28a-GST using ligation-independent cloning (LIC) for recombinant protein purification ^48,49^. Sequences were validated by whole plasmid sequencing service (Plasmidsaurus). All constructs and primers used in this study are listed in Table S5 and Table S6.

### RWT4 domain architecture prediction

RWT4^Ca^ protein architecture was predicted via InterPro (https://www.ebi.ac.uk/interpro)^32^. KDup was identified based on InterPro annotation and further defined by aligning the partial kinase sequence (amino acids 1-142; nucleotides 117-426) and the kinase domain (amino acids 189-240; nucleotides 567-720). The linker region was defined based on Alphafold structure modeling^50^. The cDNA and amino acid sequences of RWT4^Ca^ were aligned with Clustal Omega Multiple Sequence Alignment (MSA) server (https://www.ebi.ac.uk/jdispatcher/msa/clustalo) and dot plots were generated with the program Dotter obtained from Ubuntu repositories (https://packages.ubuntu.com/)^51^.

### Recombinant protein expression and purification

Full length RWT4^Ca^, the kinase region (K, 1-478 amino acids), and the pseudokinase region (PK, 540-914 amino acids) were expressed in *E. coli* BL21 Rosetta (DE3) strain and purified by affinity and size exclusion chromatography. Four liters of cells were grown at 37°C to OD_600_ of 0.4 before adding IPTG to a final concentration of 0.5 mM. Cells were resuspended in a lysis buffer consisting of 50 mM HEPES pH 8.5, 300 mM NaCl, 1 mM PMSF (phenylmethylsulfonyl fluoride) and 1 μg/mL DNase and lysed by a microfluidizer at 18k psi. The lysate was then clarified by centrifugation at 16000 g at 4°C for 50 min. The proteins of interest were separated from the clarified lysate by immobilized metal affinity chromatography (IMAC) and eluted with an elution buffer (50 mM HEPES at pH 8.5, 300 mM NaCl and 250 mM imidazole) on an AKTA FPLC. Eluted samples were dialyzed into a low-salt buffer (20 mM Tris-HCl, pH 8.5, 200 mM NaCl) for further affinity-based purification using 1 mL MBPTrapHP columns (Cytiva Life Science, #29048641). Target proteins were eluted by a buffer (10 mM maltose, 20 mM Tris-HCl, pH 8.5, 200 mM NaCl) and were injected to a HiLoad 16/600 Superdex 200 PG column equilibrated with 10 mM HEPES at pH 8.0 and 150 mM NaCl. Fractions corresponding to the proteins of interest were collected and visualized by Coomassie-stained SDS-PAGE, before being concentrated using a 3 kDa MWCO Amicon Ultra centrifugal filter (Millipore Sigma, #UFC9003) to appropriate concentrations for further analysis.

AvrPWT4 and VirPWT4 were expressed in *E. coli* BL21 Rosetta (DE3) and purified by affinity and size exclusion chromatography. Their expression and cell lysates were prepared as described above. Cell lysates were purified by affinity-based chromatography using 1mL GSTrap HP columns (Cytiva Life Science, #17528201) equilibrated with a binding buffer 140 mM NaCl, 2.7 mM KCl, 10 mM Na_2_HPO_4_, 1.8 mM KH_2_PO_4_, pH 8.0. Target proteins were eluted with elution buffer (50 mM Tris-HCl, 10 mM reduced glutathione, pH 8.0) and dialyzed to a buffer condition (10 mM Tris-HCl pH 8.0, 150 mM NaCl, 0.5 mM EDTA, 1 mM DTT) optimized for TEV protease cleavage (ProTEV Plus, Promega, V6101). Samples were adjusted to a final volume of 10 mL with 100U TEV protease and incubated at 4°C for overnight. After the incubation, samples were loaded into a GSTrap HP column (Cytiva Life Science, #17528201), and the flow-through was collected for further size exclusion chromatography (HiLoad 16/600 Superdex 200 PG column) as described above. Purified proteins were visualized by Coomassie-stained SDS-PAGE and concentrated using a 3 kDa MWCO Amicon Ultra centrifugal filter (Millipore Sigma, #UFC9003).

### Kinase activity and phosphatase assays

Kinase reactions were performed according to Lin and their colleagues with minor modifications ^52^. Kinase reactions were performed in a kinase buffer consisting of 20 mM Tris-HCl (pH 7.5), 10 mM MgCl_2_, 1 mM CaCl_2_, 1 mM dithiothreitol, 100 µM ATP, with additional 10 μCi of [^32^P]γATP (Revvity, #BLU002A100UC). Two µg of recombinant protein was mixed with the mentioned buffer and reactions were incubated at 30°C for 30 min and stopped with 3x Laemmli sample buffer. Radioactive samples were separated on SDS-PAGE gels and visualized by autoradiography. For trans-phosphorylation assays, RWT4^Ca^ and PWT4 were mixed at 1:1 ratio and incubated on ice for 20 min before the mentioned kinase reaction. For phosphatase treatments, 2 µg of recombinant protein was mixed with 400U lambda protein phosphatase (NEB, #P0753S) at 30°C for 30 min before separated by an SDS-PAGE. The phosphorylation status of RWT4 was visualized by a Western blot using an Phospho-Threonine/Tyrosine Antibody (Cell signaling technology, #9381) at 1:2000 dilution in a TBST buffer for an overnight incubation at 4°C. The antibody was detected by an Immun-Star Goat Anti-Rabbit (GAR)-HRP Conjugate (Bio-Rad #1705046) and protein signals were detected on the immunoblot using SuperSignal™ West Pico PLUS Chemiluminescent Substrate (ThermoFisher Scientific #34580) and visualized by ChemiDoc™ Touch Gel Imaging System (Bio-Rad #1708370).

### Mass spectrometry (MS)

Kinase reactions using 6 µg of RWT4^Ca^ and AvrPWT4 were conducted as described above. Samples were treated with freshly prepared 10 mM dithiothreitol and 55 mM Iodoacetamide for reduction and alkylation. Samples were digested by using trypsin at 10 ng/µL at 37°C overnight and peptides were harvested by with 50% ACN/0.1% FA (formic acid, Sigma #06473), lyophilized and resuspended with 0.1% FA. Samples were submitted to the Genomics Center Proteomics Core at the University of California, Davis. Reverse-phased Liquid Chromatography (LC) was performed on an EASY-nLC II HPLC (Thermofisher Scientific). The digested peptides were desalted by ZipTip with 0.6 µL C18 resin (ZTC18S096, Millipore) before separated by a 75-µm × 150-mm C18 100A 3-units reverse phased column at a flow rate 300 nL/min. The gradient elution employed buffer A (0.1% formic acid) and buffer B (100% acetonitrile) over 60 min, transitioning from 5% to 35% buffer B over 45 min, followed by a 35% to 80% buffer B gradient over 5 min, holding at 80% buffer B for 2 min, and finally returning to 5% buffer B over 2 min, maintaining at 5% buffer B for 6 min before the next sample injection.

Mass spectra were collected on an Orbitrap Exploris Mass Spectrometer (ThermoFisher Scientifics) with one full scan (300–1,600 m/z, R = 60,000 at 200 m/z) at a target of 3x 10^6^ ions, followed by data-independent MS/MS scans with higher-energy collisional dissociation (HCD) detected in the Orbitrap (R = 15,000 at 200 m/z). Results were analyzed by using the DIA-NN software package ^53^. The phosphorylated peptides were mapped to the sequence of RWT4^Ca^ and AvrPWT4. Peptide spectra were visualized by using Skyline software package ^54^.

### Bio-Layer Interferometry (BLI)

BLI experiments were conducted by using the ForteBio Octet® RED384 equipment at the UC Davis Octet® Real-Time Drug and Protein Binding Kinetics Unit. Analyte proteins (His-RWT4^Ca^, MBP-His-RWT4^Ca^, MBP-His-K and MBP-His-PK) were diluted to 100nM in Octet® Kinetics Buffer (SARTORIUS, #18-1105) for immobilized to Octet® Ni-NTA (NTA) Biosensors (SARTORIUS, #18-5101). Ligand proteins (AvrPWT4 and VirPWT4, without tags, or GST) were prepared at 10 µM in Octet® Kinetics Buffer as a stock solution for a 1.5x serial dilution to generate concentration gradient. Wavelength measurement for the interaction between ligand and analyte proteins followed the manufacturer’s protocol with 600 sec for association and 600 sec for dissociation ^55^.

### Yeast-two hybrid (Y2H)

To validate RWT4 and PWT4 interactions, *Rwt4^Ca^* and its variant proteins were cloned into the prey vector pGADT7 (Takara, #630442) while *AvrPwt4* and *VirPwt4* were cloned into the bait vector pGBKT7 (Takara, #630443). These constructs were co-transformed into the yeast strain AH109 following the manufacturer’s instructions from Frozen-EZ Yeast Transformation II™ (Zymo Research). Yeast transformation was tested by dilution plating cells from a starting concentration of OD_600_ 5.0 on double drop out (DDO) plates (SD-2: SD/-Trp/-Leu), while interactions were confirmed through quadruple drop out (QDO) selection (SD-4: SD/-Trp/-Leu/-His/-Ade/3-AT). 3-AT (3-amino-1,2,4-triazole) was used at 10 mM to reduce background HIS activation. The transformed yeast cells were incubated at 30°C for two to five days before imaging.

For protein expression in yeast cells, five mL of yeast cells containing the genes of interest were grown at 30°C for overnight in SD-2 and harvested by centrifuging at 5000 rpm for 10 min. Cell pellets were resuspended in a lysis buffer (150 mM NaCl, 10 mM HEPES, pH 7.5) with Pierce™ Protease Inhibitor (ThermoFisher Scientific #A32965) as suggested by the manual. Resuspended cells were mixed with acid-washed glass beads (Sigma #G8772) and lysed by sonication for 5 min. Soluble proteins were separated by centrifuging at 13000 rpm for 10 min, mixed with 3x laemmli buffer, and resolved by SDS-PAGE gels and transferred to PVDF membranes (Bio-Rad #1620177) for western blotting. Monoclonal ANTI-FLAG® M2-Peroxidase (HRP) antibody was used at 1:2000 (Millipore Sigma, A8592) dilution in a TBST buffer and incubated at 4°C overnight. Protein signals were detected by using SuperSignal™ West Pico PLUS Chemiluminescent Substrate (ThermoFisher Scientific #34580) and visualized by ChemiDoc™ Touch Gel Imaging System (Bio-Rad).

### Co-immunoprecipitation assays

To confirm the *in-planta* interaction of RWT4^Ca^ and AvrPWT4, we performed *Agrobacterium tumefaciens*-mediated transient expression in *Nicotiana benthamiana*. *Rwt4^Ca^* was cloned to a dexamethasone-inducible system (pTA7001, ^56^) with a C-terminal 3xFLAG tag, while *AvrPwt4* and *VirPwt4* were cloned to pTA7001 vector with a GFP tag. The cloned constructs were transformed into the *A. tumefaciens* C58C1 strain and infiltrated into four-week *N. benthamiana* at an OD_600_ = 0.3 for each construct. Twenty-four hours post-infiltration, 30 µM dexamethasone solution containing 0.01% Triton X-100 was applied to the leaf surface and two grams of leaf tissue was harvested 16 hours after dexamethasone application.

For immunoprecipitation, leaf tissue was ground in liquid nitrogen and resuspended in 2 mL IP buffer (50mM Tris-HCl ph7.5, 150mM NaCl, 0.1% Triton, 0.2% NP-40) containing 1× complete protease inhibitor (Thermo Fisher Scientific #A32963). Samples were centrifuged at 14000 rpm for 10 min to remove tissue debris. The supernatant was incubated with 20 µL of ChromoTek GFP-Trap® Agarose (Proteintech, # AB_2631357) at 4°C for 1 hour. Samples were washed five times with high stringency wash buffer (50mM Tris-HCl ph7.5, 300mM NaCl, 0.1% Triton, 0.2% NP-40), resuspended in 3x laemmli buffer, resolved by SDS-PAGE and transferred to PVDF membranes (Bio-Rad # 1620177) for immunoblotting. Monoclonal ANTI-FLAG® M2-Peroxidase (HRP) antibody (Millipore Sigma, A8592) or GFP Antibody, HRP (Miltenyi Biotec, # 130-091-833) were diluted at 1:2000 in a TBST buffer and incubated at 4°C for overnight. Protein signals were detected by using SuperSignal™ West Pico PLUS Chemiluminescent Substrate (ThermoFisher Scientific #34580) and visualized by ChemiDoc™ Touch Gel Imaging System (Bio-Rad).

### Rice protoplast transformation and cell viability assay

Protoplasts were prepared according to ^57^, with modifications. Seeds of the rice cultivar Kitaake (*Oryza sativa*) were surface sterilized and grown on ½ MS media at 25°C in dark for 12 days before transitioning to a photoperiod of 12 h light (approximately 150 μmol m^-2^ s^-1^) and of 12 h darkness for two days. Rice plants were cut into approximately 1 mm strips and immediately transferred into 0.6 M mannitol for 10 min in the dark.

The rice strips were then incubated with an enzyme solution (1.5% Cellulase RS, 0.75% Macerozyme R-10, 0.6 M mannitol, 10 mM MES at pH 5.7, 10 mM CaCl_2_ and 0.1% BSA) for 5 h in the dark with gentle shaking at 40 rpm. The enzymatic digestion was stopped by adding an equal volume of W5 solution (154 mM NaCl, 125 mM CaCl_2_, 5 mM KCl and 2 mM MES at pH 5.7), followed by a gentle shaking by hands for 10 seconds. Protoplasts were released and washed through 40 μm nylon mesh into round-bottom tubes three times using W5 solution. The washed protoplasts were harvested by centrifugation at 380 g for 10 min, and pellets were resuspended in MMG solution (0.4 M mannitol, 15 mM MgCl_2,_ and 4 mM MES at pH 5.7) at a concentration of 6 × 10^6^ cells/mL, determined using a hematocytometer.

For protoplast transfection, 10 µg of plasmid DNA was mixed with 100 µL of the protoplasts and 100 µL freshly prepared PEG solution [40% (W/V) PEG 4000; Sigma, 0.2 M mannitol, and 0.1 M CaCl_2_]. The mixed samples were incubated in the dark for 20 min at room temperature. After incubation, 2 mL W5 solution was added to stop the transfection, and protoplasts were pelleted by centrifugation at 380 g for 10 min. The protoplasts were resuspended in 1 mL fresh W5 solution and cultured in the dark for 16 h for cell viability assay or 4 h for detecting protein expression.

Protein expression from protoplasts were assayed by pull-down assay and Western blot. Four individual transfection events (equals to 2.4x10^6^ cells) were collected and harvested by centrifugation at 380 g for 5 min. Cell pellets were resuspended with a lysis buffer (150 mM NaCl, 10 mM HEPES, pH 7.5) containing Pierce™ Protease Inhibitor (ThermoFisher Scientific #A32965) as suggested by the manual. Cells were lysed by strong vortex for 5 min and mixed with 20 µL of Pierce™ Anti-DYKDDDDK Magnetic Agarose (Thermo Scientific™, #A36797). The mixtures were incubated at 4°C for two hours and wash for three times with the lysis buffer as suggested by the manual. Samples were mixed with 3x laemmli buffer and resolved by SDS-PAGE gels and transferred to PVDF membranes (Bio-Rad # 1620177) for immune blotting. Monoclonal ANTI-FLAG® M2-Peroxidase (HRP) antibody (Millipore Sigma, A8592) was used at 1:2000 dilution in a TBST buffer and incubated at 4°C for overnight. Protein signals were detected by using SuperSignal™ West Pico PLUS Chemiluminescent Substrate (ThermoFisher Scientific #34580) and visualized by ChemiDoc™ Touch Gel Imaging System (Bio-Rad).

### Cell viability assay

Protoplast cell viability was assayed using the CellTiter-Glo® 2.0 Cell Viability Assay kit (Promega, #G9241) as instructed by the manual and by ^58^. In brief, 100 µL of protoplasts resuspended in W5 were mixed with 100 µL of the CellTiter-Glo reagent in a 96-well white assay plate. The samples were mixed at orbital shaker for five minutes and then incubated in the dark for 10 min at room temperature prior to recording the luminescent signal. Luminescence was measured using the TriStar LB 941 Multimode Microplate Reader and MikroWin 2000 software (Berthold).

### RNA extraction and qPCR

Total RNA was extracted from rice protoplasts seven hours after transfection by using the PicoPure™ RNA Isolation Kit (ThermoFisher Scientific, #KIT0204) with on-column DNase treatment as per manual instructed. First-strand cDNA was synthesized using M-MLV Reverse Transcriptase (Promega, #M1701) with an oligo d(T) primer at 25 µg/mL. The transcripts of defense marker genes were quantified by using iTaq™ Universal SYBR® Green One-Step Kit (BIO-RAD, #1725150) with 40 cycles of 1 s at 95°C and 30 s at 60°C.

## Data analyses and availability

For protoplast viability assays, at least five independent transfection events were used as biological replicates, and the presented results are representative of at least three independent experiments. For qPCR of defense gene activation, four independent transfection events were used. Statistical differences were detected using one way ANOVA coupled with a Tukey HSD post-hoc test, α= 0.05. Protein assays including kinase activity tests, western blot, co-immunoprecipitation, and yeast-two hybrid were independently repeated at least twice with similar results. All materials are available from the corresponding author upon request. Raw data underlying each figure are available at https://doi.org/10.5281/zenodo.11087676.

## Acknowledgements

This research is supported by grants from the United States National Science Foundation (1937855) and the United States Department of Agriculture (2020-67013-32577) to GC and the United States-Israel Binational Science Foundation (BSF, 2019654) to TF and the Swiss National Science Foundation (310030_204165) to BK. MS analysis was carried out at the Genomics Center Proteomics Core at the University of California, Davis. The kinase assay using [^32^P]γATP was conducted in the facility established by Dr. Yen-Wen Kuo, Department of Plant Pathology at the University of California, Davis. BLI experiments were performed in the Cortopassi Lab with technical assistance from Dr. Alexey Tomilov at the University of California, Davis. Rice Kitaake seeds were provided by Dr. Pamela Ronald, Department of Plant Pathology at the University of California, Davis. The pUC19::Zm*UBI* plasmid was provided by Dr. Jorge Dubcovsky from the Department of Plant Science, University of California, Davis. The pET28s::GST vector was provided by Dr. Nitzan Shabek, Department of Plant Biology, University of California, Davis.

## Author contributions

Y-CS and GC designed the study with assistance from TZ and BK. SA performed wheat inoculation experiments, YL performed yeast-two hybrid, ZB identified KDup, SB assisted with molecular cloning, *Agrobacterium*-mediated expression in *N. benthamiana* and western blotting under the guidance of Y-CS and GC. Y-CS performed all other experiments and data analyses. Y-CS and GC wrote the original draft and all authors reviewed and edited the manuscript.

## Ethics declarations

The authors declare no competing interests.

