## Supplementary Figures S1-S9 for "Direct binding of a fungal effector by the wheat RWT4 tandem kinase activates defense"

**Supplementary materials:**

**Supplementary tables S1-S6**

Table S1 Reaction of *Triticum monococcum* and *T. aestivum* accessions to Br48 and Br48+*AvrPwt4*
Table S2 Binding kinetics (Kd) of His-RWT4 to AvrPWT4 and VirPWT4
Table S3 Binding kinetics (Kd) of His-MBP-RWT4, His-MBP-KD, and His-MBP-PKD to AvrPWT4
Table S4 *Rwt4* alleles from different wheat reference genomes and the blast resistance phenotypes
Table S5 Constructs used in this study
Table S6 Primers used in this study

**Supplementary Figures S1-S9**

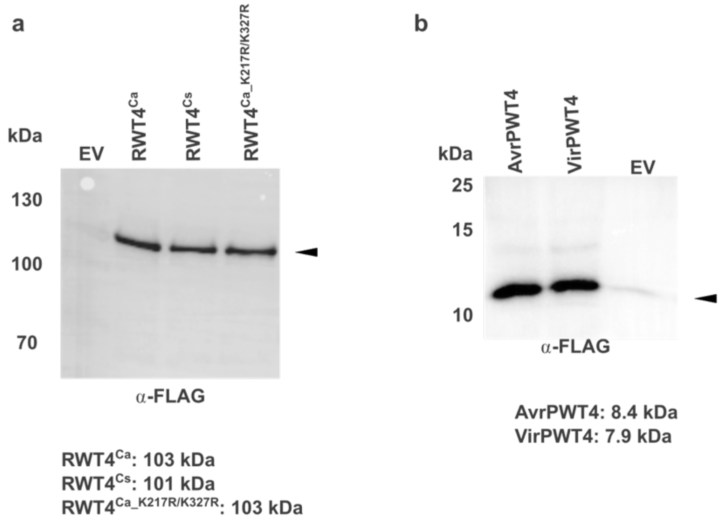


**Figure S1. RWT4 and PWT4 expression from rice protoplasts. (a)** Western blot showing expression of RWT4$^{Ca}$, RWT4$^{Cs}$, and RWT4$^{Ca_K217R/K327R}$ as well as **(b)** AvrPWT4 and VirPWT4 from rice protoplasts. 2.4x$10^6$ protoplast cells were harvested and lysed for pull-down assay using Pierce™ Anti-DYKDDDDK Magnetic Agarose (Thermo Scientific™, #A36797). Samples were resolved by SDS-PAGE gels and transferred to PVDF membranes for western blotting using ANTI-FLAG® M2-Peroxidase (HRP) antibody (1:2000 dilution, Millipore Sigma, A8592). Expected molecular weight of the proteins are indicated.

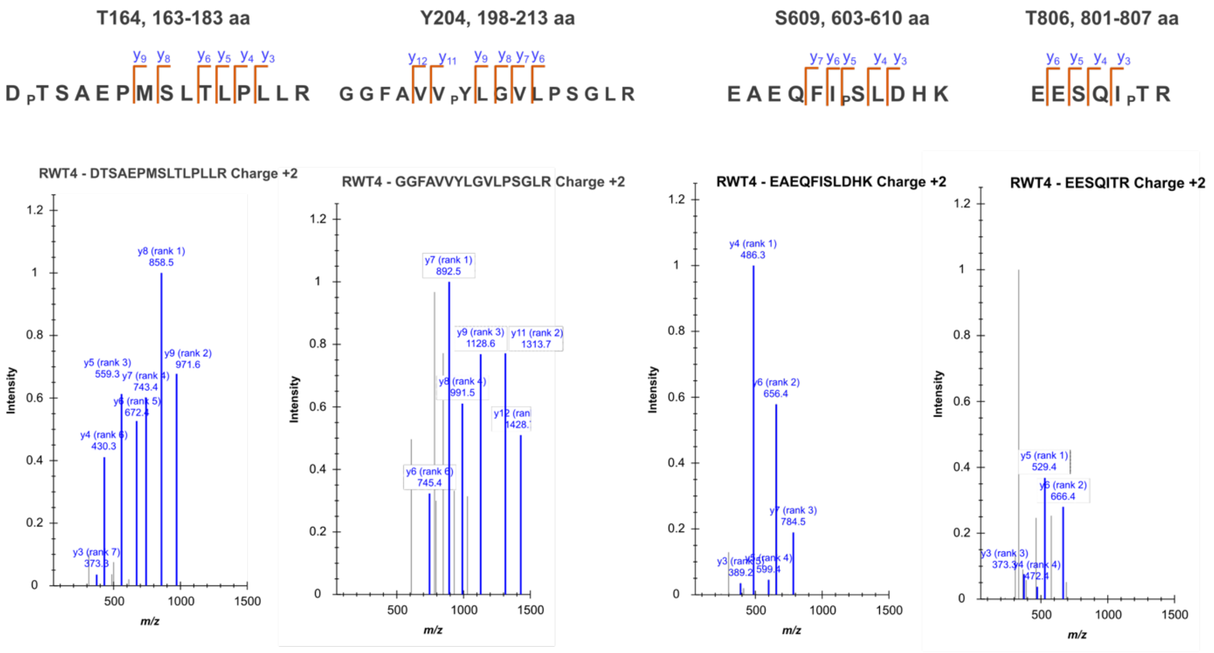


**Figure S2. Mapping RWT4$^{Ca}$ phosphorylation sites upon AvrPWT4 recognition.** Recombinant proteins were incubated in kinase reaction buffer (RWT4 +/- AvrPWT4) and subjected to mass spectrometry to identify phosphorylated residues. Four phosphorylated residues were identified, with T164, Y104 in the kinase region and S609, S803 in the pseudokinase domain. Peptide spectra were visualized by using the Skyline software package.

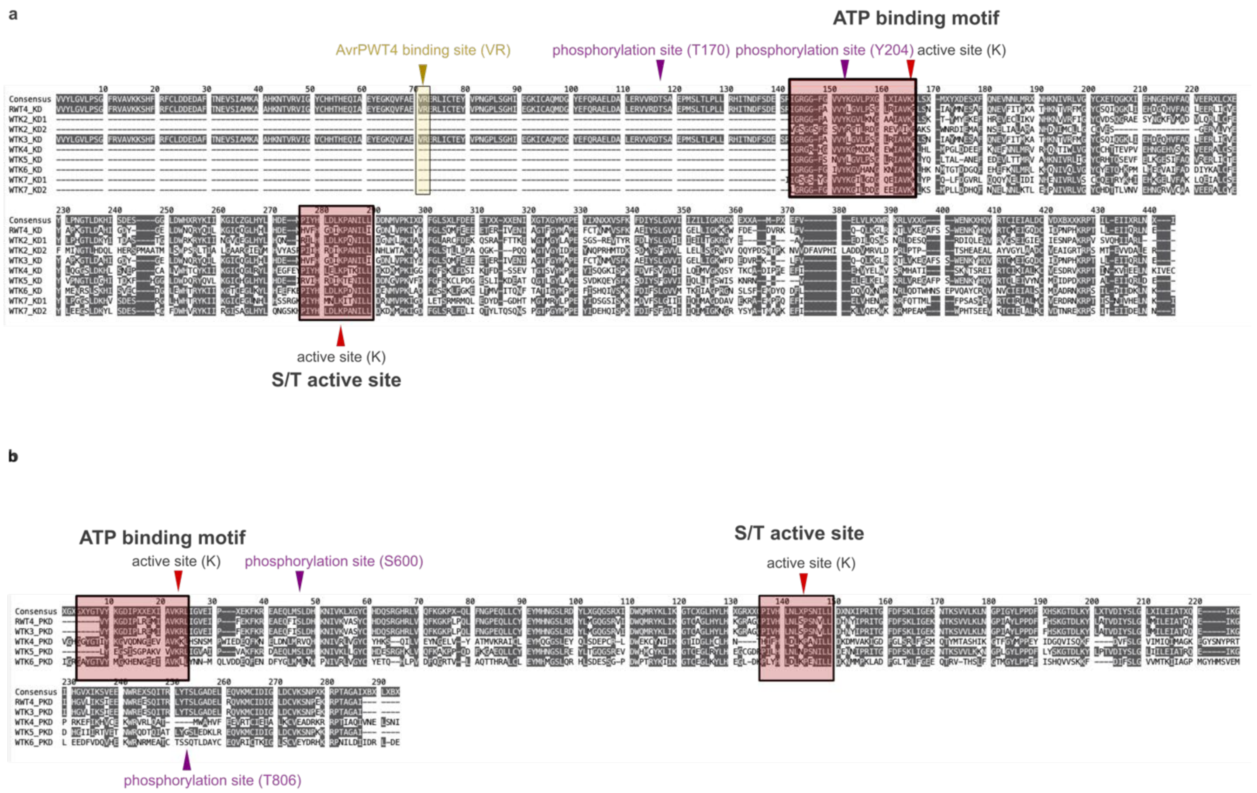


**Figure S3. Sequence alignment of TKP LRR_8B subfamilies. (a)** Kinase and **(b)** pseudokinase from TKPs within the LRR_8B kinase subfamily, including RWT4, WTK2, WTK3, WTK4, WTK5, WTK6, and WTK7-Tm. Red boxes indicate ATP binding motif or S/T active site predicted via InterPro. The lysine (K) within kinase's ATP binding loop and S/T active site is conserved across all tested TKPs, while the lysine within the pseudokinase's S/T active site is not conserved. The four identified RWT4$^{Ca}$ phosphorylation sites are indicated in purple. The identified AvrPWT4 binding site is highlighted yellow.

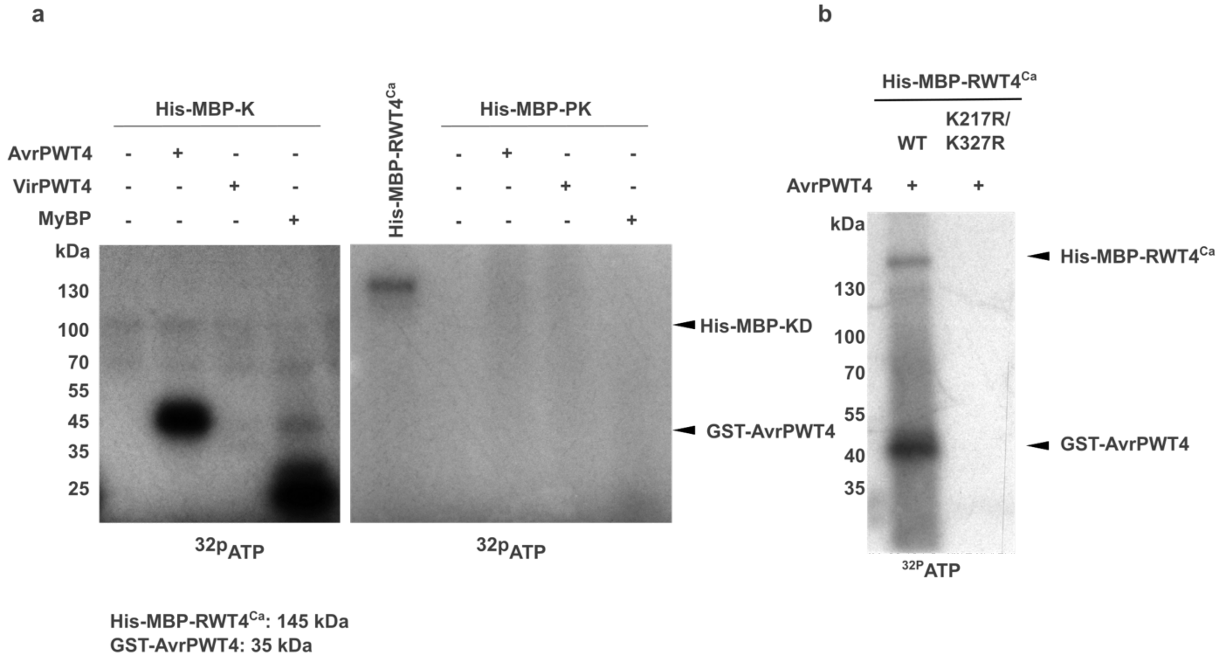


**Figure S4. Full-length RWT4, as well as the kinase region only, can transphosphorylate AvrPWT4.** Transphosphorylation activity was assayed by incubating 6 µg of **(a)** His-MBP-Kinase (K, amino acids 1-478), His-MBP-Pseudokinase (PK, amino acids 540-914), or **(b)** His-MBP- $RWT4^{Ca\_K217R/K327R}$ with AvrPWT4, VirPWT4, or MyBP at room temperature for 20 min. After the incubation, kinase reaction was conducted by mixing samples with [γ-32P]ATP and were then resolved by running SDS-PAGE. Phosphorylation of the substrate proteins were visualized via autoradiography.

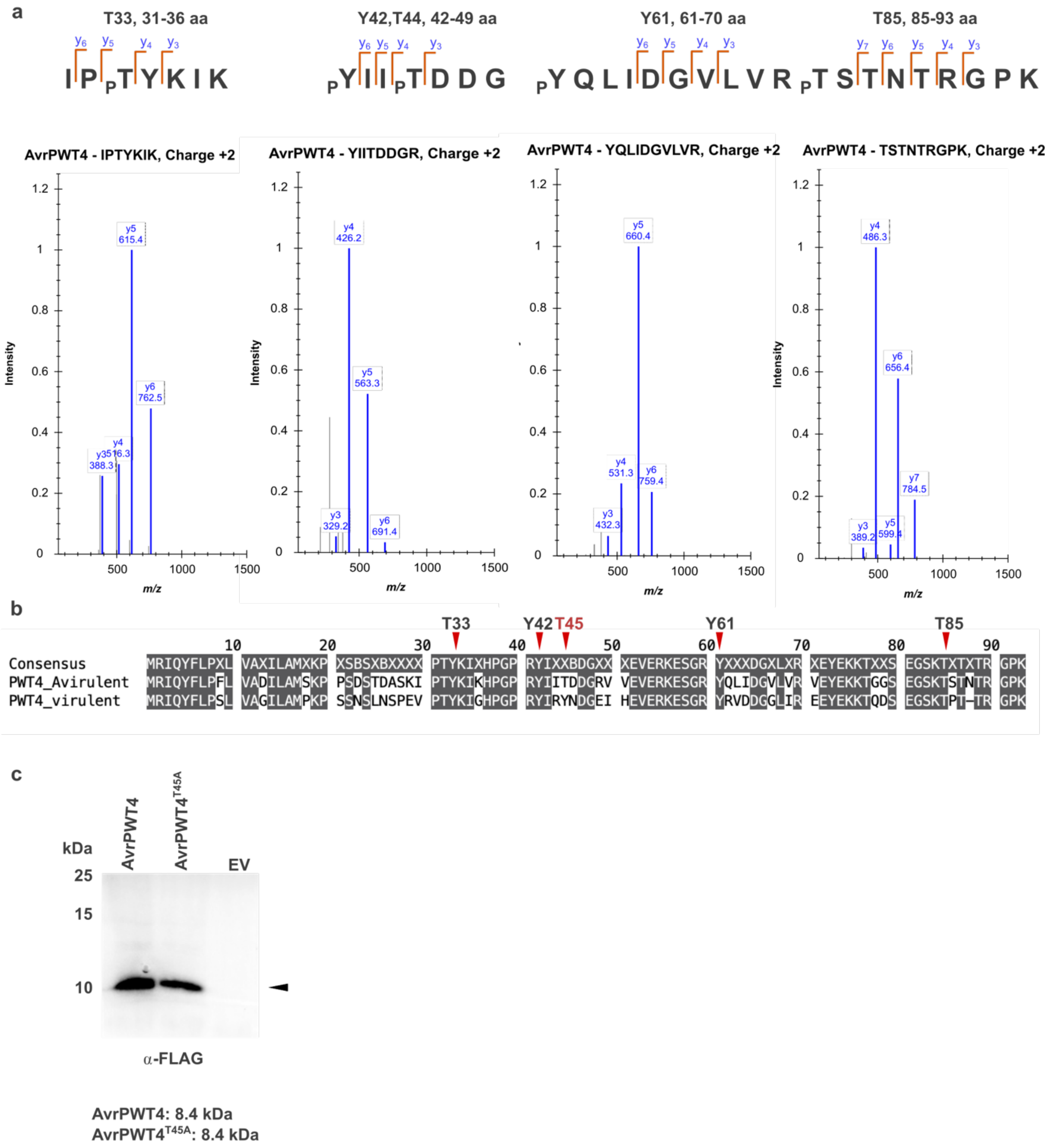


**Figure S5. Mapping sites on AvrPWT4 that are directly phosphorylated by RWT4$^{Ca}$. (a)** Recombinant proteins were incubated in a kinase reaction buffer and subjected to mass spectrometry to identify phosphorylated residues. Five AvrPWT4 phosphorylated residues were identified: T33, Y42, T45, Y61, and T85. Peptide spectra of the identified phosphorylated peptide from AvrPWT4. Peptide spectra were visualized by using the Skyline software package. **(b)** Sequence alignment between AvrPWT4 and VirPWT4. The identified phosphorylation sites are indicated by arrows and the unique T45 AvrPWT4 phosphorylation site is labelled in red. **(c)** Western blot showing protein expression of AvrPWT4 and AvrPWT4$^{T45A}$ in rice protoplasts. 2.4x10$^{6}$ protoplast cells were harvested and lysed for pull-down assay using Pierce™ Anti-DYKDDDDK Magnetic Agarose (Thermo Scientific™, #A36797). Samples were resolved by SDS-PAGE gels and transferred to PVDF membranes for immune blotting using ANTI-FLAG® M2-Peroxidase (HRP) antibody (1:2000 dilution, Millipore Sigma, A8592). Expected molecular weight of the proteins are indicated.

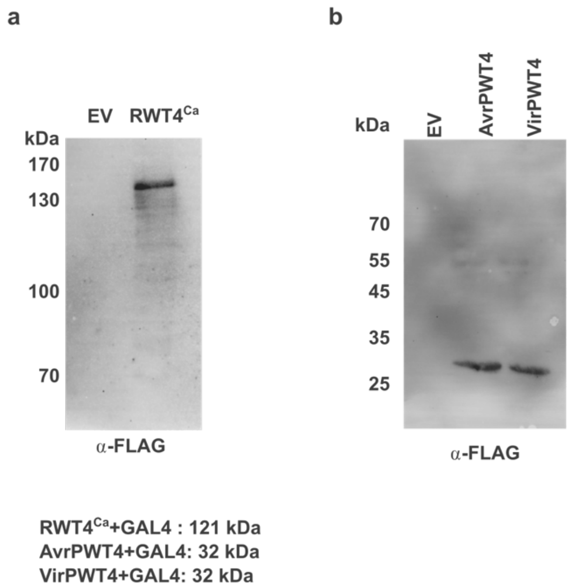


**Figure S6. Expression of RWT4$^{Ca}$, AvrPWT4 and VirPWT4 from yeast.** Cells were harvested from a 5mL overnight culture. Samples were resolved by SDS-PAGE and transferred to PVDF membranes for western blotting using ANTI-FLAG® M2-Peroxidase (HRP) antibody (1:2000 dilution, Millipore Sigma, A8592). Expected molecular weights of the proteins are indicated.

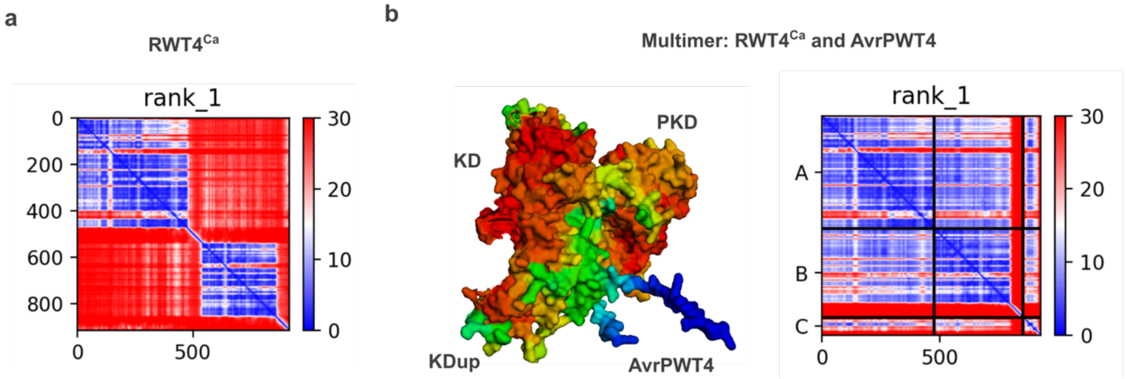


**Figure S7. Confidence values for the Alphafold2 prediction of the RWT4$^{Ca}$ structure and Alphafold-multimer modeling of RWT4$^{Ca}$ and AvrPWT4.** (a) Predicted Aligned Error (PAE) plot of the predicted RWT4$^{Ca}$ structure. (b) Alphafold-multimer modeling of RWT4$^{Ca}$ and AvrPWT4 colored by pLDDT scores (left) and its PAE plot (right).

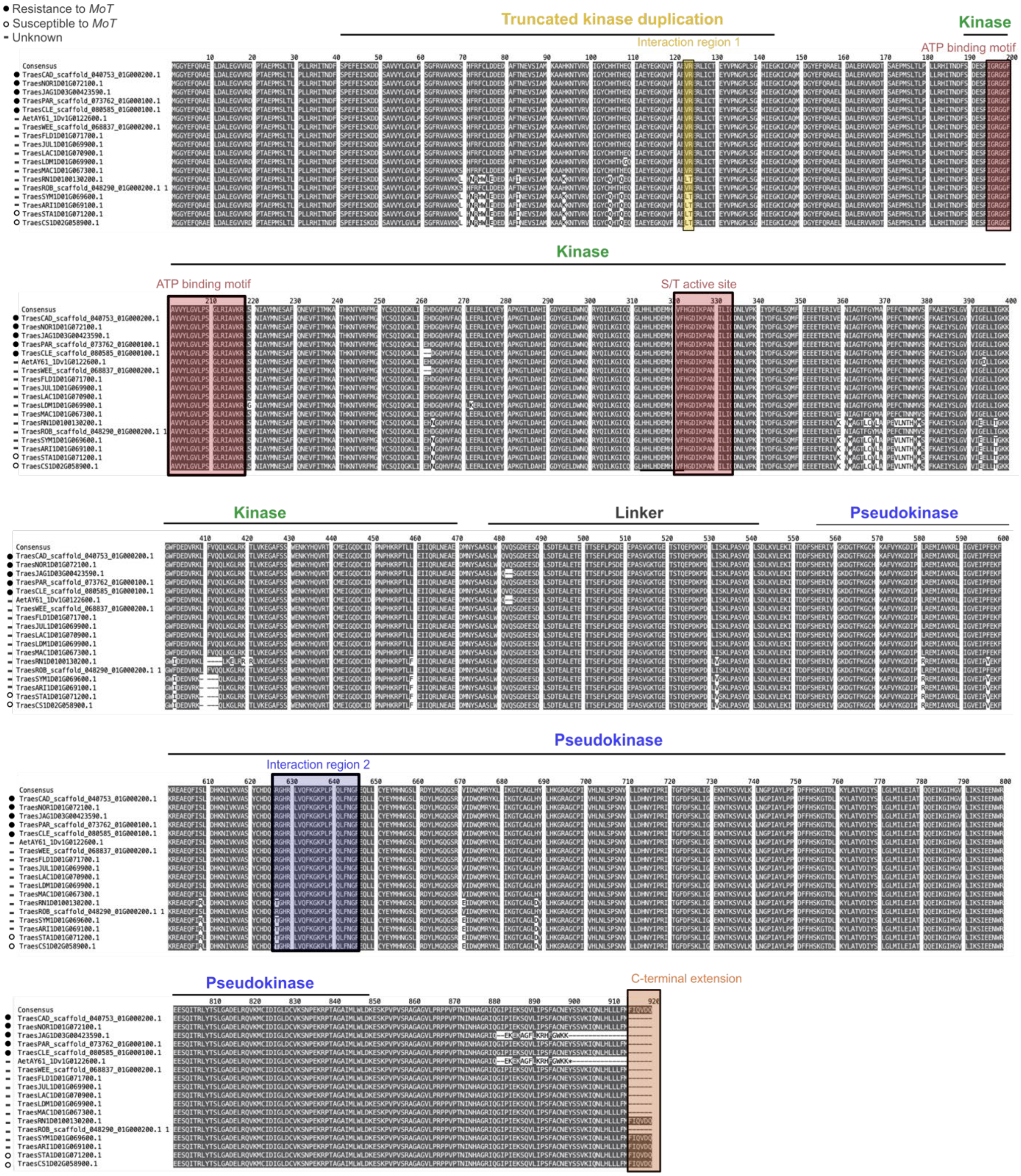


**Figure S8. Amino acid sequence alignment of RWT4 from different alleles.** RWT4 sequence identity and their source cultivars are listed in Table S4. Integrated truncated kinase duplication (KDup), kinase domain, pseudokinase domain and linker are highlighted. The yellow box indicates the interaction region I identified from Fig 5b. Red boxes indicate the ATP binding motif and kinase site predicted via InterPro [29]. The C-terminal extension region is labeled orange. Known resistant alleles are labeled with a solid black circle, and susceptible alleles with an open circle.

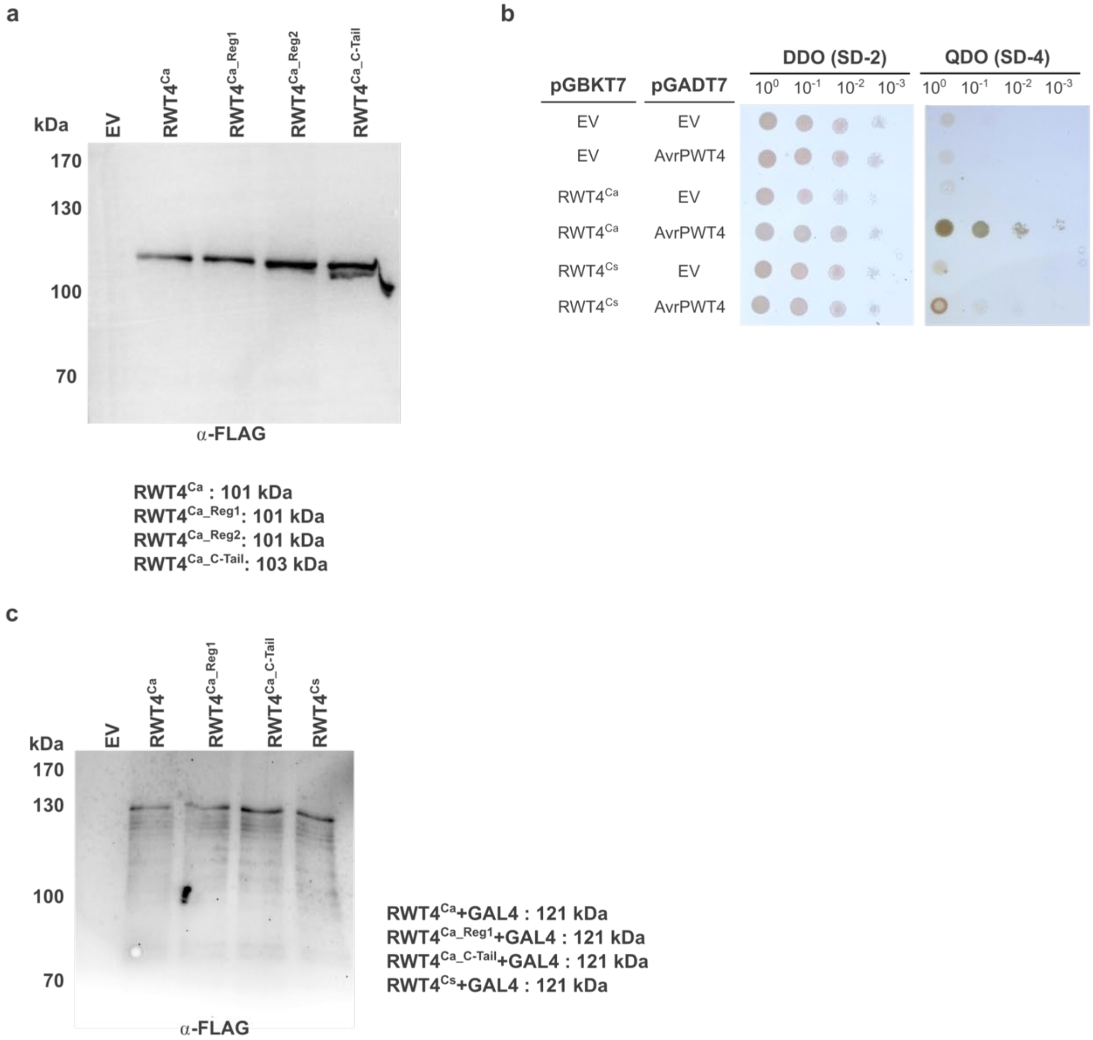


**Figure S9. Expression of RWT4$^{Ca}$ variant proteins from yeast and rice protoplasts. (a)** Expression of RWT4 variant proteins including RWT4$^{Ca_Reg1}$, RWT4$^{Ca_Reg2}$, RWT4$^{Ca_+Ctail}$ from rice protoplasts. 2.4x10$^{6}$ protoplast cells were harvested and lysed for pull-down assay using Pierce™ Anti-DYKDDDDK Magnetic Agarose (Thermo Scientific™, #A36797). Samples were resolved by SDS-PAGE gels and transferred to PVDF membranes for immune blotting using ANTI-FLAG® M2-Peroxidase (HRP) antibody (1:2000 dilution, Millipore Sigma, A8592). Expected molecular weight of the proteins are indicated. **(b)** Yeast two-hybrid showing the RWT4$^{Ca}$ interaction with AvrPWT4. The susceptible allele RWT4$^{Cs}$ exhibited weak interaction with AvrPWT4. Yeast colony growth on DDO (SD-Leu/-Trp) media indicates the presence of the two plasmids and the growth on QDO/X/A (SD-Ade/-His/-Leu/-Trp/3-amino-1,2,4-triazole) media indicates interaction. Serial dilutions from cell suspensions of a single yeast colony expressing the bait and prey plasmids are shown and represent the strength of interaction. Empty vectors (EV) of pGBKT7 and pGADT7 were used as negative controls. The presented results are representative of two independent replicates. **(c)** Expression of RWT4 variant proteins in yeast. Cells were harvested from a 5mL overnight culture. Samples were resolved by SDS-PAGE gels and transferred to PVDF membranes for immune blotting using ANTI-FLAG® M2-Peroxidase (HRP) antibody (1:2000 dilution, Millipore Sigma, A8592). Expected molecular weights of the proteins are indicated.
